# The multifunctional scaffold protein Small ovary couples piRNA-guided transposon recognition to nuclear RNA decay and heterochromatin formation

**DOI:** 10.64898/2026.08.12.744517

**Authors:** Zsuzsanna Földi, Melinda Bence, Zsanett Takács, Erika Gábor, Aladár Pettkó-Szandtner, Róbert Tóth, Anna Poscher, Viktor Vedelek, Rita Sinka, Viktor Honti, Ildikó Unk, Zsolt Czimmerer, Miklós Erdélyi, Ferenc Jankovics

## Abstract

The piRNA pathway maintains genome integrity by silencing transposons cotranscriptionally in the nucleus through recognition of nascent transposon RNAs and recruitment of endogenous transcriptional and chromatin-level repressive mechanisms to transposon loci. However, the molecular link between the SFiNX complex, which recognizes nascent transposon RNA, and downstream effector complexes has remained elusive. Here, we demonstrate that the Small ovary (Sov) protein mediates this connection. By mapping the functional activities of its structural elements, we reveal that Sov contributes to transposon silencing through two distinct molecular mechanisms. First, Sov specifically directs nascent transposon transcripts toward nuclear RNA exosome-mediated degradation by physically interacting with the RNA decay factor TEsup1. Second, Sov directly binds the heterochromatin protein HP1a via multiple conserved motifs and undergoes phase separation, facilitating heterochromatin formation and genome-wide gene repression. Genetic analyzes of *sov* mutants reveal that these functions are separable: RNA-mediated transcriptional silencing is essential for piRNA pathway activity, while phase separation-dependent heterochromatin regulation is critical for stable transposon repression. We propose that Sov acts as a molecular scaffold in piRNA-guided transposon silencing, integrating transposon recognition with cotranscriptional RNA decay and chromatin-based regulatory pathways.

## INTRODUCTION

Mobile genetic elements, or transposons, are widespread components of the genomes of living organisms. The remobilization of active transposons in the germline poses a constant threat to the integrity of a species’ genome; therefore, organisms have evolved diverse strategies to prevent their activity. This process is best characterized in the *Drosophila* ovary, where transposon expression is selectively controlled by the piRNA pathway, in which short, 25–30 nt long single-stranded piRNAs play a central role.^1^ In *Drosophila*, the piRNAs are associated with Piwi-type proteins, such as Piwi, Ago3, and Aub and silence transposon expression at both transcriptional and post-transcriptional levels. The post-transcriptional silencing takes place in the cytoplasm of germ cells, where the piRNA-bound Aub and Ago3 proteins cleave complementary transcripts via their slicer activity (Post-Transcriptional Gene Silencing, PTGS).^2,3^ Concurrently, in both somatic and germline cells, Piwi-bound piRNAs translocate to the nucleus to suppress transposon transcription at their genomic loci (Transcriptional Gene Silencing, TGS).^4–7^

The initial step of the transcriptional silencing pathway is the recognition of a complementary sequence on the nascent transposon transcript by the Piwi-bound piRNA. Subsequently, the target-engaged Piwi/piRNA complex promotes the assembly of the Piwi-Asterix/Gtsf1-Maelstrom complex, which functions as a molecular platform that enables the recruitment of an additional downstream complex, the SFiNX (also known as Pandas/PICTS/PPNP) complex, to the nascent transcript.^8–12^ The dimeric SFiNX complex, composed of Panx, Nxf2, Nxt1, and Lc8, couples target recognition to the distinct downstream processes of transposon silencing.^13,14^ The Nxf2 subunit of the SFiNX complex binds to the target-engaged, activated Piwi, while the N-terminus of the Panx subunit undergoes Piwi-dependent SUMOylation and recruits the multi-zinc-finger protein, Small Ovary (Sov).^11,15^ The association of Sov with the SFiNX complex is essential for the initiation of all subsequent downstream processes in transcriptional transposon silencing.^15,16^ By an unknown mechanism, Sov activates endogenous silencing pathways at the transposon loci suppressing their expression at two levels of gene regulation: RNA metabolism and localized heterochromatinization.^16–18^

For RNA-level silencing, the core transcription termination machinery is actively repurposed as a functional arm of the piRNA system. The SFiNX complex recruits proteins that stalls RNA polymerase II at the transposon loci, facilitating transcriptional termination and subsequent exosomal degradation of the nascent transposon RNAs.^16,19,20^ According to the current model of the RNA-level transposon silencing, PNUTS directs Protein Phosphatase 1 (PP1) to the transcriptional apparatus to dephosphorylate the elongation factor Spt5. This modification promotes the recruitment of Pcf11, to the termination zone, where phase-separated Pcf11/Spt5 condensates form. These condensates decelerate RNAPII and Senataxin resolves R-loop (RNA–DNA) hybrids at transcriptional pause sites to promote the release of Pol II from the transposon DNA template. The fate of these terminated transposon transcripts is determined by TEsup1/CG31510 and TEsup2/CG7065, which physically interact with scaffold proteins of various nuclear exosome adaptor complexes.^20^ The recruitment of the NEXT, TRAMP and PAXT complexes directs these transcripts toward nuclear exosomes for degradation.

Chromatin-level silencing relies on the coordinated action of various chromatin-modifying enzymes and regulatory factors, including histone methyltransferases, demethylases, deacetylases, and chromatin remodelers.^21^ Upon target recognition, the SFiNX complex facilitates the recruitment of the SetDB1/Wde histone methyltransferase complex, leading to the deposition of the repressive H3K9me3 histone mark. This modification is indispensable for the recruitment of HP1a and the subsequent establishment of heterochromatin at Piwi-targeted transposon loci.^6,8,22^ In parallel, histone marks associated with transcriptionally active chromatin are systematically removed: the histone demethylase Lsd1 eliminates H3K4 mono- and dimethylation to impede transcription initiation, while the histone deacetylase Rpd3 drives chromatin compaction and transcriptional repression.^23–25^ Furthermore, chromatin accessibility is modulated by the linker histone H1.^26,27^ Notably, Sov may also play a direct structural role in heterochromatin formation, as evidenced by the fact that its loss leads to genome-wide heterochromatin depletion.^28^

A striking feature of Piwi-induced transposon repression is that several of its constituent processes, such as the recognition of nascent transposon transcripts by the SFiNX complex, RNAPII stalling by Pcf11/Spt5, and heterochromatic repression by HP1a, occur within phase-separated nuclear domains.^14,19,29^ The formation of these membraneless nuclear compartments is driven by liquid–liquid phase separation (LLPS), which is mediated and maintained by multivalent interactions between intrinsically disordered proteins or regions (IDPs/IDRs).^30^ The association of Sov with the SFiNX complex is indispensable for transposon silencing.^15^ Sov has been proposed to function as a critical scaffold to orchestrate the recruitment of both the transcriptional inhibitory machinery and heterochromatin-forming factors to transposon loci.^16^ However, the specific molecular activities of Sov required for transposon silencing or heterochromatin regulation remain elusive.

Here, we show that the Sov protein directly interacts with HP1a at multiple binding sites and identify specific intrinsically disordered regions (IDRs) that drive its phase separation and are required for heterochromatin formation. We demonstrate that Sov mediates transcriptional silencing through both a SFiNX-dependent, RNA-level inhibitory process by promoting RNA decay and a broader, phase-separation-dependent, genome-wide heterochromatin regulation. Importantly, our findings reveal that transcriptional inhibition of nascent transcripts is essential for the subsequent formation of a repressive heterochromatic environment at transposon loci.

## RESULTS

### Sov associates with distinct nuclear domains

Although the SFiNX complex plays a key role in transposon recognition, transcriptional repression, and initiating heterochromatin modifications, it remains evenly distributed throughout the nucleoplasm.^9^ In contrast, its interactor Sov, a heterochromatin-promoting factor required for transposon silencing, accumulates in heterochromatin.^17,18^ This discrepancy suggests that Sov functions in transcriptional repression and chromatin regulation in spatially separated nuclear domains. To test this, we re-examined the subnuclear localization of Sov in germ cell nuclei using high-resolution microscopy after coexpression of the full-length Sov tagged with EGFP (Sov(FL):EGFP) and HP1a:RFP in egg chambers (Figure1B). High-resolution imaging revealed three distinct localization patterns for Sov(FL):EGFP: (1) a diffuse fraction distributed throughout the nucleoplasm, (2) accumulation within HP1a:RFP-positive heterochromatic foci, and (3) small, non-heterochromatic foci lacking HP1a:RFP (FigureS1). As the distribution of the non-heterochromatic foci resembled the branched morphology of the nucleolus, we coexpressed Sov(FL):EGFP with the nucleolar marker Fibrillarin:RFP (Fib:RFP) and confirmed that these foci associate with nucleolar compartments (Figure1C).

**Figure 1.**
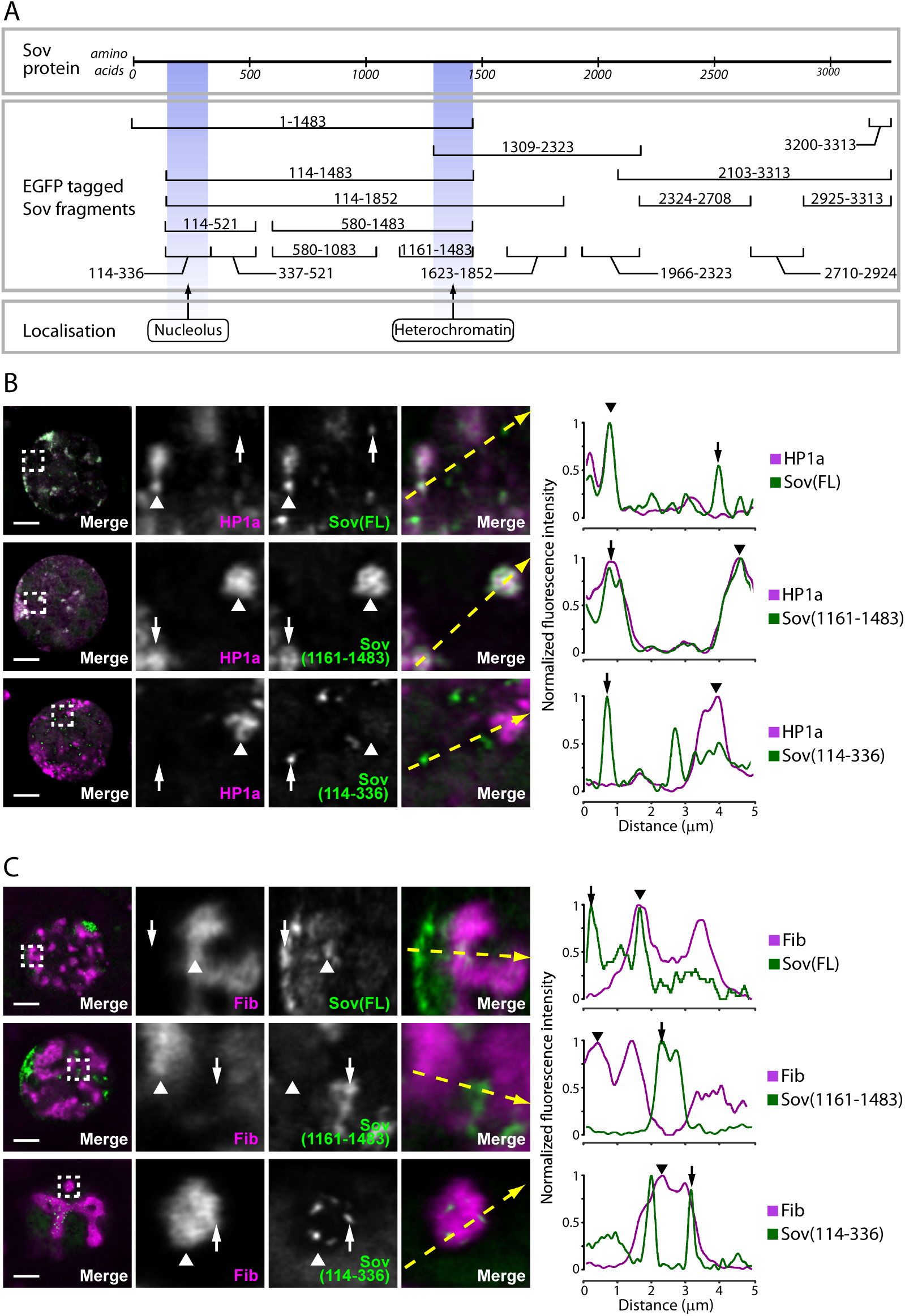
Sov is localised at multiple nuclear foci. **(A)** Schematic diagram of Sov protein structure and fragments tested for subnuclear localization. Amino acid coordinates of each fragment are indicated. Colored boxes indicate the specific regions responsible for nucleolar and heterochromatic accumulation. **(B, C)** Colocalization of EGFP-tagged FL Sov and Sov fragments with HP1a:RFP (B) and Fib:RFP (C). Confocal microscopy images display germ cell nuclei. Germ cell-specific expression of the EGFP-tagged Sov protein fragments is under the control of the *nos*-Gal4 driver. Insets show higher-magnification views of the dashed boxes. Right panels display fluorescence intensity profiles quantified along the yellow dashed arrows. Arrowheads and arrows indicate corresponding points between images and diagrams. Scale bars represent 5 µm.

To map the regions responsible for this multi-domain targeting, we divided Sov into overlapping fragments. Each of the fragments was fused to NLS-EGFP and coexpressed with HP1a:RFP (Figure1A, FigureS1). The mapping revealed that Sov(1161–1483) selectively mediates localization to HP1a:RFP-positive heterochromatin, whereas Sov(114–336) directs localization to Fib:RFP-positive nucleolar domains (Figure1B,C). Although these two fragments localized to non-overlapping domains but both perfectly colocalized with full-length Sov (FigureS1B). Together, these results demonstrate that Sov localizes to all major nuclear compartments implicated in transposon silencing, suggesting its participation in both heterochromatin-dependent and -independent silencing mechanisms. Moreover, its nucleolar localization implies a function in nucleolus-associated processes.

### Sov forms phase separated condensates

Given that both RNA-dependent transposon silencing and heterochromatin formation are associated with IDR-mediated phase separation, and that the nucleolus itself represents a phase-separated compartment, we examined whether Sov contains IDRs capable of driving this process. Using IUPRED and PONDR prediction algorithms, we identified a long continuous disordered region in the N-terminal half of the protein, Sov(114–1483), and two shorter disordered segments: one located centrally between predicted Zn-finger domains, Sov(1623–1852), and another at the extreme C-terminal end, Sov(3200–3313) (Figure2A).

**Figure 2.**
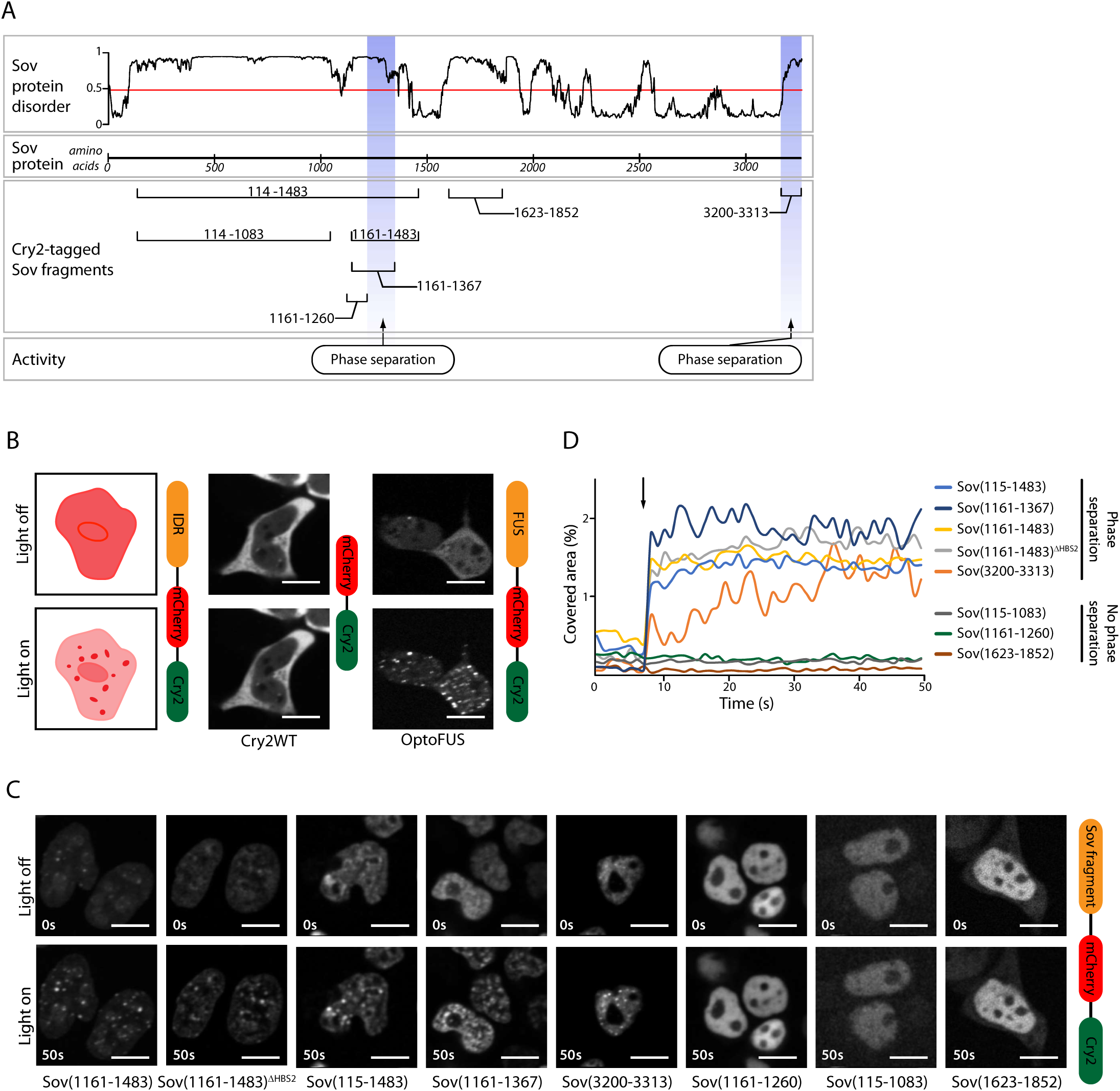
Intrinsically disordered regions of Sov drive phase separation. **(A)** Schematic representation of Sov protein disorder profile (top) predicted by IUPRED3 and mapping of Cry2:mCherry-tagged Sov fragments used in the optoDroplet assay.^78^ Blue boxes indicate regions exhibiting phase separation activity. **(B)** Schematic diagram of the optogenetic system (left) and representative confocal images showing light-induced clustering. Cry2WT serves as a negative control (middle), while OptoFUS as a positive control for light-induced phase separation (right). Top row shows cells before blue light exposure (Light off); bottom row shows condensation after illumination (Light on). Scale bars represent 10 µm. **(C)** Representative time-lapse images of various Cry2:mCherry-tagged Sov fragments before (Light off, top row) and after (Light on, bottom row) blue light activation. Scale bars represent 10 µm. **(D)** Quantification of the kinetics of light-induced phase separation for the indicated Sov fragments (TableS1). The graph plots the area covered by the condensates (in % of the nuclear area, n>19 nuclei). The blue light illumination pulse is indicated by an arrow at 7s.

Since IDRs typically display high thermotolerance, we performed a thermostability assay to confirm the *in silico* prediction. Analysis of two purified Sov fragments, Sov(157-816) and Sov(819-1299), encompassing 83% of the predicted N-terminal disordered region demonstrated *in vitro* that the fragments maintain solubility upon extended heat treatment strongly supporting their unstructured nature (FigureS2A).

To functionally test the phase separation potential of these regions, we applied the optoDroplet assay, an *in vivo* optogenetic approach that relies on light-induced oligomerization of Cryptochrome 2 (Cry2). This structural change drives the clustering of the Cry2-coupled protein fragments, which nucleate droplet formation only when the fragment itself possesses phase-separating capacity. (Figure2B).^31^ Sov IDRs were fused to Cry2:mCherry and expressed in HEK293T cells. Upon light-induced clustering, droplet formation was observed for two disordered regions of Sov: the 1369 amino acid-long N-terminal, Sov(114-1483) and the extreme C-terminal, Sov(3200–3313) (Figure2C, TableS1). Further mapping of this N-terminal segment narrowed the droplet forming activity to a 226 amino acid-long segment, Sov(1161-1483). Truncating this fragment to 106 amino acids abolished droplet formation, indicating that the Sov(1260–1367) region is responsible for driving phase separation (Figure2A). In summary, we identified two regions within Sov capable of mediating phase separation: Sov(1260–1367) and Sov(3200–3313). Both regions contain conserved stretches and are enriched in serine and arginine residues, a feature commonly observed in phase-separated proteins (FigureS2).^32^

We next investigated the kinetics of droplet assembly by quantifying the area fraction occupied by droplets (Figure2D). The protein fragments containing the Sov(1260–1367) region promoted rapid droplet formation, whereas Sov(3200–3313) promoted slower droplet growth. Analysis of droplet motility revealed that Sov(1260–1367) droplets were static. As this protein region largely overlaps with the heterochromatin-associating segment Sov(1309– 1483), these immobile aggregates likely associate with heterochromatin. In contrast, Sov(3200–3313) droplets were mobile, suggesting they can diffuse within the nucleoplasm. These findings imply that the two phase-separating regions may mediate distinct functions of Sov.

### Sov interacts with the SFiNX complex and HP1a through multiple conserved binding motifs

The association of the SFiNX complex with transposon mRNA leads to heterochromatin formation at the transposon locus, implying an interaction between the recognition and heterochromatin-forming complexes.^8^ To investigate the potential role of Sov in this process, we sought to identify its direct binding partners. To this end, we performed a candidate-based yeast two-hybrid (Y2H) screen including components of the SFiNX complex (Panx, Piwi, Nxf2) as well as the general heterochromatin factor HP1a. To map the interaction sites within Sov, we divided the protein into eight fragments and tested their binding to the selected interaction partners (Figure3A, FigureS3).

**Figure 3.**
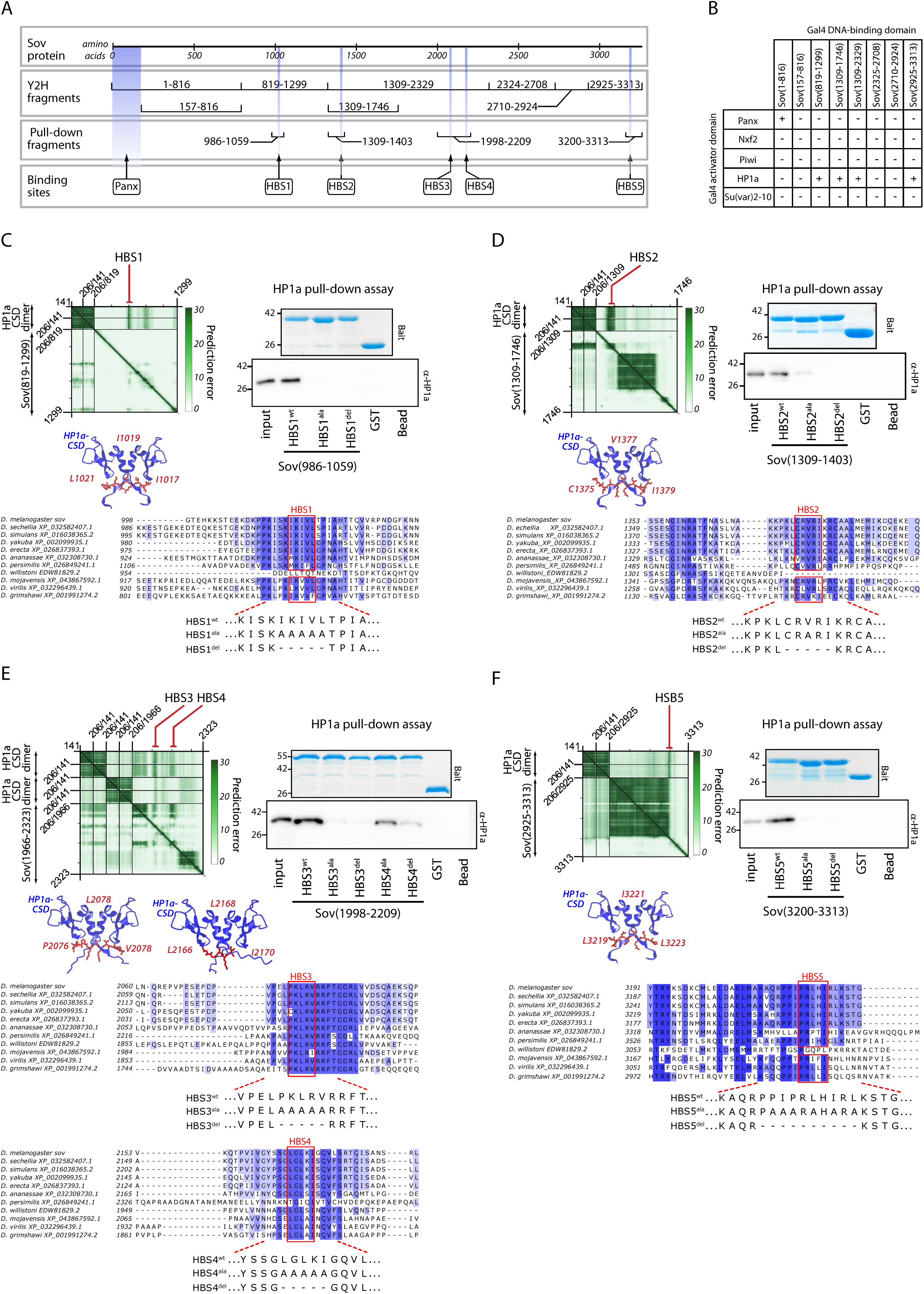
The Sov protein contains multiple distinct HP1a-binding sites. **(A)** Schematic overview of the Sov protein (top), mapping the fragments used for Y2H assays (middle) and GST pull-down assays (bottom). Amino acid coordinates of each fragment are indicated. The positions of the identified Panx-binding region (Panx) and five HP1a-binding sites (HBS1 to HBS5) are indicated at the bottom. **(B)** Y2H interaction matrix between various Gal4 DNA-binding domain (BD)-fused Sov fragments and Gal4 activator domain (AD)-fused proteins (Panx, Nxf2, Piwi, HP1a, and Su(var)2-10). Plus (+) signs indicate interaction; minus (–) signs denote lack of interaction. **(C–F)** Structural prediction and biochemical validation of individual HP1a-binding sites: HBS1 in (C), HBS2 in (D), HBS3 and HBS4 in (E), and HBS5 in (F). Left panels: AlphaFold-Multimer prediction alignment error (PAE) matrices for the indicated Sov fragments complexed with the HP1a chromo shadow domain (CSD) dimer (top), alongside the corresponding structural models (middle). Critical interacting amino acid residues of the conserved motifs (red) are highlighted. Right panels: Representative Western blots of GST pull-down assays verifying the physical interactions, using purified recombinant His-HP1a as a ligand and purified GST-tagged wild-type (wt), alanine-substituted (ala), or deletion (del) mutant Sov fragments as bait. Bait proteins were monitored in the pull-down samples via Coomassie blue staining, whereas His-HP1a was detected using an anti-HP1a antibody. GST-only and bead-only samples served as negative controls. Bottom panels: Multiple sequence alignments showing the evolutionary conservation of HBSs across representative *Drosophila* species. Shading intensity corresponds to the degree of amino acid conservation. Amino acid sequences of wild-type (wt), alanine-substituted (ala), or deletion (del) mutant HBSs are indicated.

Consistent with previous findings, Sov was found to bind Panx, and this interaction was mapped to the N-terminal region encompassing amino acids 1–157 (Figure3B, FigureS3).^15^ Furthermore, Y2H assays revealed that three regions of Sov are capable of binding HP1a (Figure3B). To further examine these interactions, we generated AlphaFold3 models to predict direct contacts between HP1a dimers and the respective Sov fragments. Based on the predictions, five putative HP1a binding sites (HBS1 to HBS5) were identified within Sov (Figure3C-F). The binding sites were located within extended IDRs, each corresponding to a variant of the PxVxL pseudopalindromic HP1a-binding consensus short linear motif and showing evolutionary conservation (Figure3C-F).^33–35^ Recombinant protein fragments containing these HBS regions were expressed as GST fusions in bacteria and purified for *in vitro* binding assays with purified His-tagged HP1a (Figure3C-F). Because HBS3 and HBS4 are in close proximity, they were examined together within a single protein fragment (Figure3E). GST pulldown assays showed that all tested fragments containing predicted HBS motifs were able to bind HP1a. For HBS1, HBS2, and HBS5, alanine-substitution or deletion of the PxVxL motif abolished HP1a binding, indicating specific interaction (Figure3C,D,F, FigureS3). In the fragment containing both HBS3 and HBS4, mutation of HBS3 nearly eliminated binding, whereas mutation of HBS4 caused only a modest reduction (Figure3E). This suggests that HBS3 mediates stronger interaction, while HBS4 contributes weakly to HP1a binding *in vitro*.

Y2H and pulldown experiments indicate a robust physical interaction with multiple binding sites between Sov and HP1a, which may engage Sov in different regulatory processes. Interestingly, HBS2 is located within the region of Sov required for both heterochromatin association and phase separation. Therefore, we examined whether HP1a binding, mediated by HBS2 is required for Sov’s phase-separation capacity. Mutation of the HP1a binding site did not impair phase separation in the optoDroplet assay, indicating that Sov retains this ability independently of HP1a interaction (Figure2C,D, TableS1).

### Recruitment of Sov to nascent RNA induces gene silencing independently of heterochromatin formation or phase separation

It has been demonstrated that Sov, when recruited by the SFiNX complex to nascent transposon RNA, initiates the transcription termination and heterochromatin formation at the targeted locus.^15,16,19^ In the previous experiments, we demonstrated that Sov establishes direct physical interactions with components of both the SFiNX complex and the heterochromatin-modifying machinery. Furthermore, Sov exhibits the capacity for liquid-liquid phase separation (LLPS), a property shared by SFiNX, HP1a, and Pcf11/Spt5, which may be mechanistically linked to transposon silencing.

To identify the specific molecular requirements for Sov-mediated RNA-based transcriptional inhibition, we applied the λN/BoxB tethering assay.^22^ In this approach, the λN-tagged Sov fragment is tethered to BoxB RNA motifs placed at the 3′ end of a constitutively expressed EGFP reporter transcript, allowing to exert an effect on the reporter transcription (Figure4A).^36^ Sov was divided into fragments, which were specifically expressed in germ cells as λN-tagged constructs (Figure4B,C). EGFP reporter gene expression was assessed at the mRNA level via RT-qPCR and at the protein level via fluorescence microscopy. This analysis identified Sov(337–521) capable of inducing RNA-based suppression (Figure4B,C). Quantitative PCR confirmed that repression occurred at the transcriptional level (Figure4D, TableS2). When a LacI-tagged Sov fragment containing this region was tethered to DNA instead of RNA via LacI/LacO interaction, it did not exhibit repressive activity, indicating that the silencing process is initiated from the nascent transcript rather than through the recognition of the encoding DNA (FigureS4B).^22^

**Figure 4.**
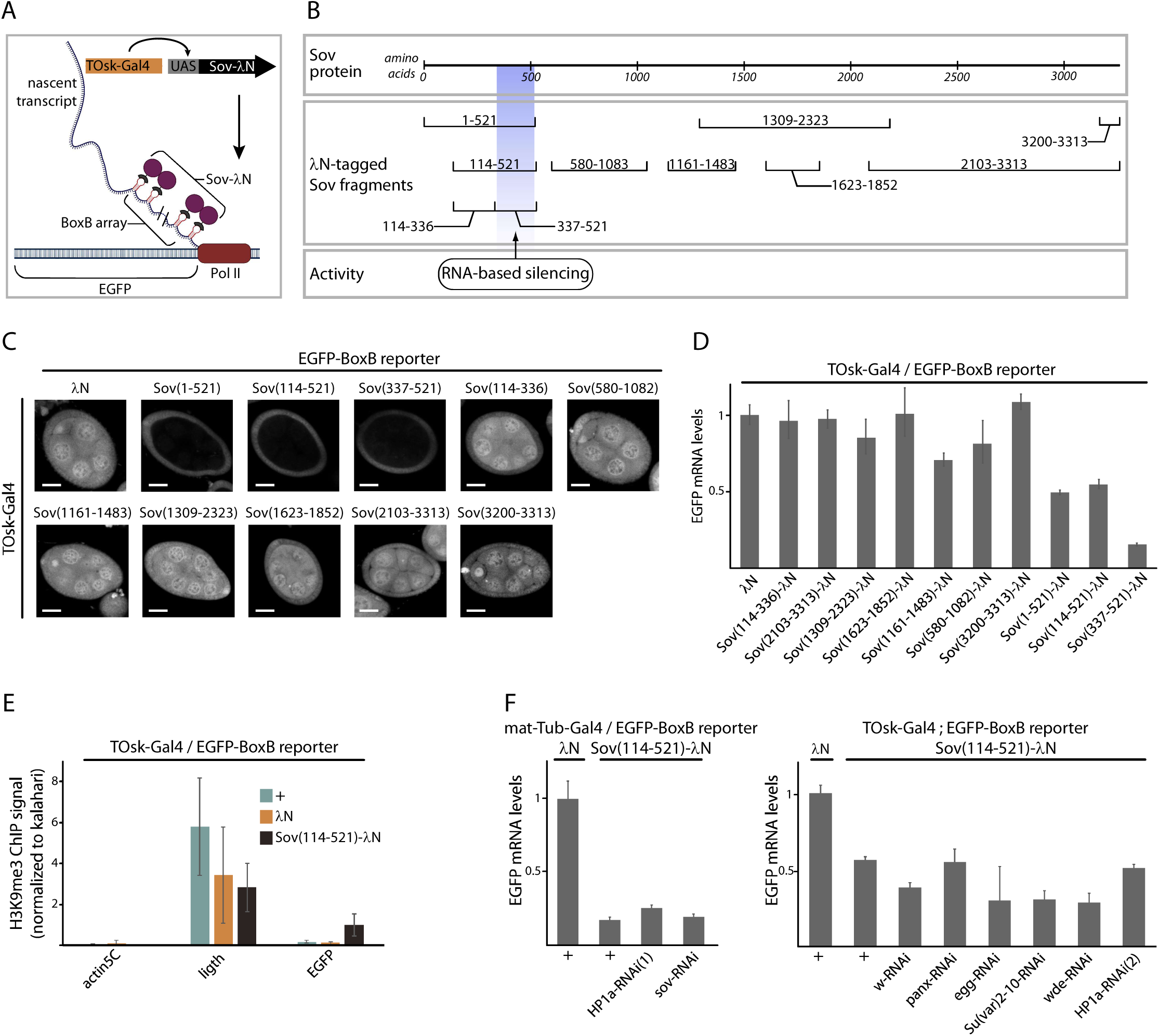
Recruitment of Sov to nascent RNA induces transcriptional silencing. **(A)** Schematic of the λN/BoxB tethering system. The expression of λN-tagged Sov protein fragments is driven by a germ line specific TOsk-Gal4 driver. **(B)** Schematic overview of the Sov protein and the λN-tagged fragments tested in the tethering assay. Amino acid coordinates of each fragment are indicated. The colored box indicates RNA-based silencing activity mapped to Sov(337–521). **(C)** Representative confocal images illustrating the silencing activity of individual Sov fragments. Egg chambers expressing the indicated λN-tagged Sov protein fragments in the germline and the EGFP-BoxB reporter in both germline and somatic cells. Germ cell-specific expression of the λN-tagged Sov protein fragments is under the control of the TOsk-Gal4driver. Scale bars represent 20 µm. **(D)** Diagram showing the quantification of the EGFP reporter mRNA levels in ovaries expressing the indicated λN-tagged Sov fragments. Data are normalized to the λN control. Error bars indicate SD. **(E)** Chromatin immunoprecipitation analysis of H3K9me3 enrichment at the EGFP reporter locus upon recruitment of the Sov(114–521) fragment measured by qPCR. Actin5C represents an active euchromatic locus, while *light* is heterochromatic locus. H3K9me3 ChIP-signal was normalised to gene desert *kalahari*. Error bars indicate SD of three biological replicates. **(F)** Quantitative RT-PCR of EGFP reporter mRNA levels in ovaries coexpressing the λN-tagged Sov(114–521) fragment and the indicated shRNAs against various heterochromatin regulators. The left panel shows the effects of shRNA constructs expressed via the mat-Tub-Gal4 driver; the right panel shows the effects when driven by the TOsk-Gal4 driver. Data are normalized to the λN control. Error bars indicate SD.

To investigate the mechanism underlying the transcriptional suppression, we examined the chromatin state of the reporter locus upon Sov tethering. Given that Sov is known to promote heterochromatin formation genome-wide, we tested for the presence of the heterochromatin-mark H3K9me3 at the reporter locus.^22^ ChIP-qPCR revealed only a low level of H3K9me3 enrichment upon Sov(114-521) tethering (Figure4E, TableS2). To further clarify the mechanism by which the Sov exerts its repressive effect, we performed a desilencing assay by expressing different shRNAs in germ cells in which Sov(114–521)-λN was tethered to BoxB-EGFP (Figure4F, FigureS4A). In line with the weak H3K9me3 signal at the EGFP reporter locus, repression mediated by Sov(114–521) did not require HP1a, Eggless (Egg), Windei (Wde) or Su(var)2-10, which are all known to contribute to heterochromatin formation at transposon loci.^22,37–40^ Furthermore, depletion of endogenous Sov had no effect on Sov(114– 521)-mediated reporter silencing, indicating that additional activities, such as heterochromatin association or phase separation, of the full-length Sov protein are not necessary for the observed transcriptional suppression (Figure4F). These results suggest that Sov-mediated RNA-based repression of the reporter occurs through a mechanism that is independent of heterochromatin formation.

### RNA-associated Sov connects transposon recognition to RNA decay

To dissect the molecular mechanism underlying RNA-mediated reporter suppression by Sov, we performed a TurboID-based proximity labeling assay coupled with quantitative mass spectrometry. In germ cells, the TurboID-tagged EGFP nanobody was coexpressed with the Sov(337–521):EGFP fragment, the shortest Sov fragment capable of mediating RNA-based transcriptional silencing. Sov interacting proteins were identified by affinity-purification and analyzed by LC–MS/MS. Quantitative analysis of the mass spectrometry data revealed that two paralogous proteins, CG31510/TEsup1 and CG7065/TEsup2, were enriched in the Sov-proximal proteome (Figure5A, TableS3). Interestingly, these two proteins have previously been identified as factors associated with RNA-binding-dependent transcriptional silencing and RNA decay, suggesting that RNA-tethered Sov silences transposons by directly interfering with RNA metabolism.^16,20^

**Figure 5.**
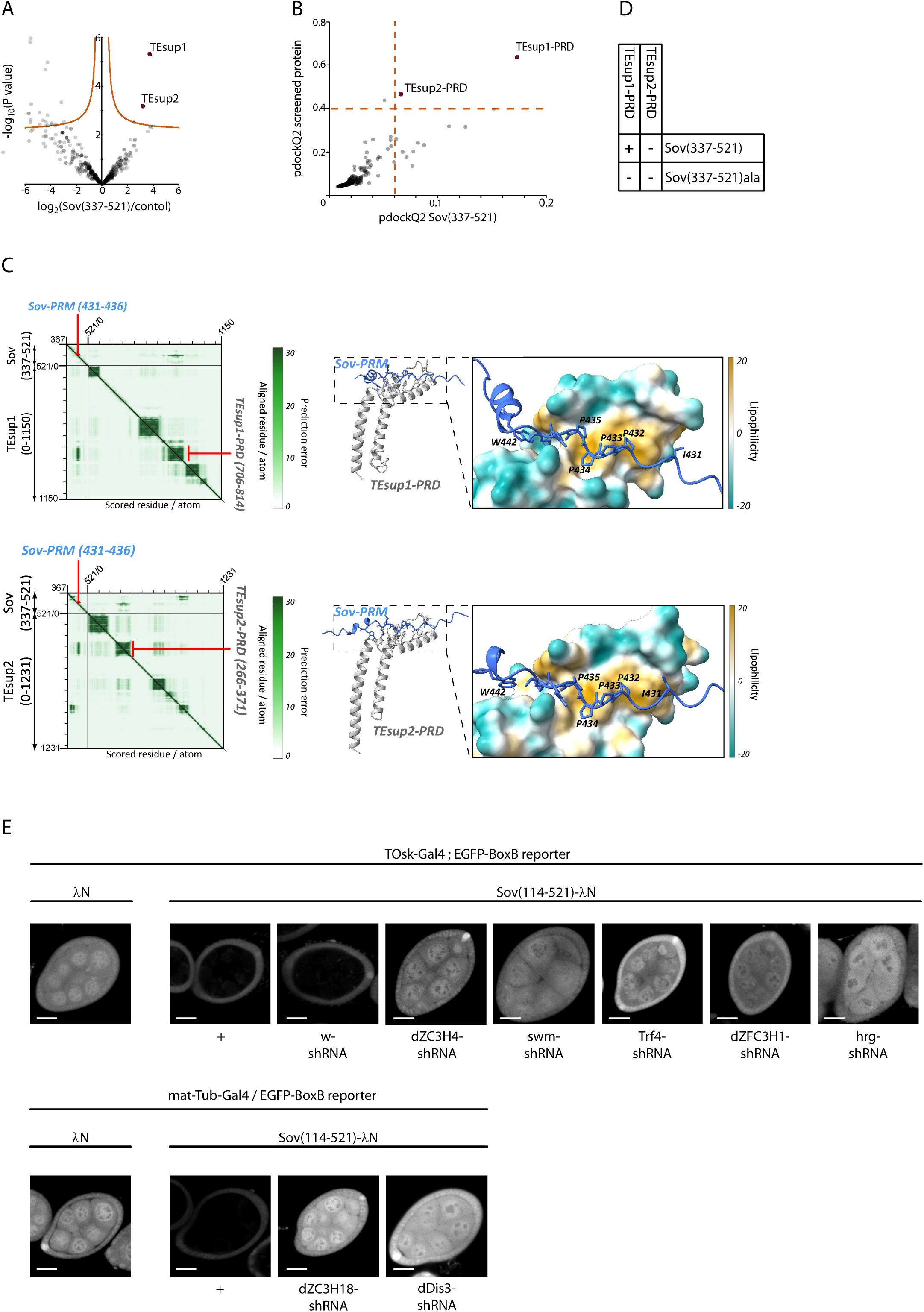
Sov-PRM Interaction with TEsup1-PRD mediates cotranscriptional suppression. **(A)** Volcano plot illustrating protein enrichment from LC-MS label free quantification analysis comparing Sov(337–521)-TurboID against the GFP-TurboID negative control (TableS3). Significantly enriched proteins are highlighted in red (False Discovery Rate, FDR < 0.05). **(B)** AlphaFold3 pairwise interaction screen between the Sov(337–521) fragment and selected candidate proteins (TableS4). Scatter plot displays the pdockQ2 scores of the screened proteins against the pdockQ2 scores of Sov(337–521), highlighting TEsup1-PRD and TEsup2-PRD as high-confidence interactors above the scoring thresholds (dashed lines). **(C)** AlphaFold3 structural prediction of the interaction complex of the Sov proline-rich motif (Sov-PRM, residues 431–436) with the TEsup1 proline-rich domain (TEsup1-PRD, residues 706–814) and with the TEsup2 proline-rich domain (TEsup2-PRD, residues 266– 371). Left: PAE plot for the Sov(337–521) and TEsup1 full-length (0–1150) and TEsup2 full-length (0-1231) interaction. Right: Structural model of the interaction interface colored by surface lipophilicity, with interacting residues of Sov-PRM (I431, P433, P434, P435, P436, and W442) highlighted. **(D)** Y2H interaction matrix validating the binding Sov-PRM/TEsup1-PRD interface. The assay scores physical interactions between the indicated target domains and either wild-type Sov(337–521) or an alanine-substituted mutant within the core PRM motif (Sov(337– 521)ala). Plus (+) signs denote interaction; minus (–) signs denote lack of interaction. **(E)** Tethering assay using the λN/BoxB reporter system to assess transcriptional silencing activity. Representative confocal micrographs show EGFP-BoxB reporter fluorescence in egg chambers. Expression of the λN-tagged Sov(114–521) fragment driven by the germline-specific mat-Tub-Gal4 or the TOsk-Gal4 drivers results in reduced EGFP reporter expression, which is de-repressed by the simultaneous shRNA-mediated silencing of the indicated RNA-decay factors. Scale bars represent 20 µm.

Next, by identifying the direct protein interactors of Sov, we sought to elucidate the suppressive mechanisms underlying transcriptional silencing. To this end, we performed an AlphaFold3-based *in silico* screen by modeling the structural interfaces between Sov(337– 521) and 76 candidate proteins previously shown to coimmunoprecipitate with Sov.^16^ In addition, we tested 46 candidate proteins involved in RNA metabolic processes that are required for transposon silencing, such as transcription termination, RNA 3’ processing, nuclear exosome targeting, and RNA decay (TableS4).^16,19^ Both protein sets included both TEsup1 and TEsup2. In total, we screened 120 full-length proteins and their 164 structured fragments for physical interactions with Sov(337–521). Consistent with our proximitome analysis, short fragments of the TEsup2 and TEsup1 proteins emerged as reliable *in silico* interactors (Figure5B, TableS4). In both proteins, an α-helical region, the so called proline recognition domain (PRD), was predicted to form high-confidence interfaces with a short proline-rich motif (PRM) within the Sov(337–521) fragment (Figure5C, FigureS5A,B).

Next, we analyzed the structural interfaces between Sov and the predicted interactor proteins *in vivo.* Y2H assays confirmed that Sov(337–521) associates *in vivo* with the PRD of TEsup1 protein through a direct physical interaction (Figure5D, FigureS5A,B). Targeted point mutations within the PRM of Sov(337–521) abolished binding to the TEsup1-PRD, demonstrating that the interaction between the PRM and the PRD recruits TEsup1 protein to Sov. However, despite the *in silico* predicted binding, Y2H analysis failed to detect a physical interaction between Sov(337–521) and TEsup2-PRD.

To investigate the functional relevance of the Sov-TEsup1 interaction, we employed the λN/BoxB-based transcription reporter assay. We tethered a Sov(114–521) fragment to the nascent BoxB-EGFP reporter transcript, which resulted in the silencing of the reporter gene expression. Simultaneously, we performed RNAi knockdown of genes involved in nuclear exosome targeting and RNA decay, previously shown to be required for transposon silencing.^20^ We observed that the depletion of the nuclear exosome subunit (CG6413/Dis3), as well as components of the TRAMP (dTRF-4), PAXT (swm, hrg, CG4294/dZFC3H1), NEXT (CG1677/ZC3H18) and Restrictor (Su(s)/ZC3H4) complexes, prevented Sov-mediated silencing of the EGFP reporter gene (Figure5E). These results suggest that RNA-bound Sov recruits downstream nuclear exosome adaptor complexes to direct nascent transcripts toward degradation.

### Sov mutants uncouple transcription and heterochromatin silencing

To assess the contribution of Sov’s distinct molecular activities to transcriptional silencing, we generated mutants targeting specific functional domains of the protein using CRISPR/Cas9-mediated genome editing (Figure6A, TableS5). Given that phase separation can influence transcriptional regulation at multiple levels, we deleted the region required for phase separation and heterochromatin localization (*sov*^3032^). To distinguish between phenotypes caused by defects in phase separation or HP1a binding, we created a mutant in which only the HP1a binding site (HBS2) was deleted (*sov^ΔHBS2^*). To dissect the genetic contribution of phase separation and HP1a interaction, we also used the *sov*^2^ allele, which contains a frameshift mutation resulting in the loss of the C-terminal phase separation domain and one of the HP1a binding site (HBS5).^18,41^ To investigate Sov’s direct role in transcriptional regulation, we deleted the region responsible for RNA-mediated silencing (*sov*^1216^). We generated two additional deletion alleles that remove large IDRs located near the RNA-based silencing domain (*sov*^1929^) or the phase-separating region (*sov*^2627^). We also included the *sov^2KE^* allele, which abolishes Sov binding to Panx.^15^ Finally, we used the *sov^def^*^1^ allele, which represents a complete deletion of the *sov* gene.^18^ We analyzed the mutant phenotypes in several functional assays, including viability, fertility, heterochromatin formation, and transcriptome profiling (Figure6B-D).

**Figure 6.**
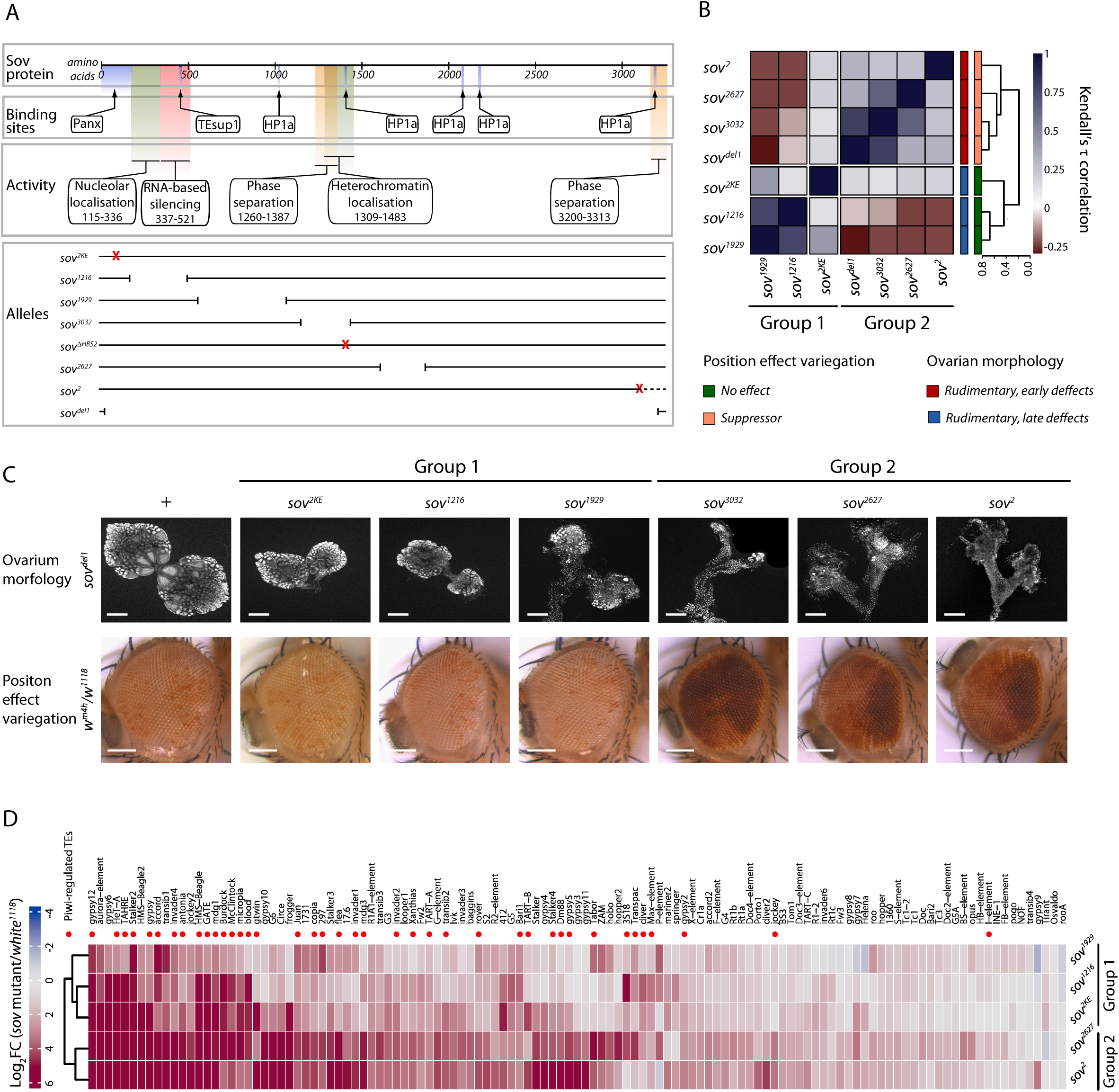
Genetic analysis reveals separable functions of Sov in transposon silencing. **(A)** Schematic overview of the Sov protein showing Panx and HP1a binding sites (top). Distinct functional activities are denoted by colored boxes, with corresponding amino acid coordinates indicated (middle). Structural alignment mapping the genomic positions of mutations corresponding to the indicated *sov* mutant alleles (bottom). Red signs indicate the positions of the frame-shift mutation in *sov*^2^, the two-amino-acid substitution in *sov^2KE^*, and the five-amino-acid deletion in *sov^ΔHBS2^*.^15,17,18^ The brackets indicate large in-frame deletions. **(B)** Correlation analysis of *sov* alleles. The heatmap displays pairwise Kendall’s τ correlation coefficients, calculated from categorical phenotypic scoring across the allelic series based on severity of the sterile phenotype (TableS6). Hierarchical clustering of the coefficients segregates the *sov* alleles into two groups. **(C)** Phenotypic profiling of *sov* alleles. Top row: Representative micrographs illustrating ovarian phenotype of the indicated alleles in trans-heterozygous over *sov^del1^*. Nuclei are labeled with DAPI, Scale bar represents 200 µm. Bottom row: Representative micrographs of adult female eyes demonstrating the dominant effect of the indicated sov alleles on PEV in the *w*^1118^/*w^m^*^4h^ background. Scale bar represents 100 µm. **(D)** Transcriptional derepression profile of transposable elements in *sov* mutant ovaries. The heatmap depicts the log_2_ fold change (log_2_FC) of TE steady-state RNA levels in *sov* mutant ovaries relative to *w*^1118^ control. All indicated alleles are trans-heterozygous over *sov^del1^* null allele.

The *sov^def^*^1^ null allele was homozygous lethal, indicating that *sov* is essential for viability. Interestingly, the *sov^ΔHBS2^* homozygous females showed no phenotype, indicating that the HP1a binding site HBS2 is not essential for the analyzed *sov* functions. All the other *sov* alleles were homozygous female-sterile, indicating that the deleted regions in the novel alleles affect germline-specific functions. The analyzed female sterile alleles were also sterile when combined with *sov^def^*^1^, confirming that in all cases the observed defects were caused by the loss of *sov* function (TableS6).

Next, we analyzed the ovarian phenotypes of transheterozygous sterile females carrying the specific *sov* alleles in combination with *sov^def^*^1^ (Figure6C). Microscopic analysis of the ovaries defined two well-separated allele groups. Group 1 included alleles impaired in RNA-mediated silencing and Panx binding (*sov*^1929^, *sov*^1216^, and *sov^2KE^*). Group 1 mutants produced mature eggs in the ovaries; however, females either failed to lay eggs or laid eggs that failed to hatch. This phenotype is characteristic of mutants affecting specific components of the SFiNX complex.^8,12,15,22,40^ Group 2 included alleles defective in phase separation, HP1a binding, or heterochromatin localization (*sov*^2^, *sov*^3032^, and *sov*^2627^). Mutants belonging to Group 2 exhibited atrophic ovaries lacking mature eggs, indicative of an early arrest in oogenesis. This phenotype is characteristic of mutations in heterochromatin regulators and suggests a general heterochromatin dysfunction in mutant germ cells.^25,39,42–47^ A comprehensive genetic complementation analysis by testing the viability and fertility of females carrying pairwise combinations of all *sov* alleles revealed that certain *sov* alleles partially or fully complemented each other (Figure6B, TableS6). Based on the penetrance of sterility in the various allele combinations, we constructed a correlation matrix that confirmed the existence of two distinct groups of the alleles, affecting either the direct transcriptional regulatory function or the heterochromatin-promoting activity of Sov. These observations suggest that Sov exerts its effects on transposon silencing through two distinct genetically separable inhibitory mechanisms: transcriptional repression and chromatin-level regulation.

This hypothesis was supported by the behavior of the mutants in the position effect variegation (PEV) assay, which was used to test the dominant effect of the *sov* mutations on heterochromatin spreading (Figure6C).^48^ Group 1 alleles failed to suppress variegation, indicating no effect on heterochromatin. This demonstrates that these Sov activities mediate transposon-specific, heterochromatin-independent silencing rather than global heterochromatin formation. Group 2 alleles, however, were dominant PEV suppressors, indicating that Sov’s heterochromatin-promoting activity requires phase separation and/or HP1a binding.

Analysis of the ovarian transcriptome of specific *sov* mutants revealed a striking difference between the two allele groups (Figure6D, FigureS6, TableS7). In Group 1 mutants (*sov^2KE^*, *sov*^1929^ and, sov^1216^) in which the transposon-specific Sov functions were impaired, mainly Piwi-regulated transposons were affected (FigureS6A). In contrast, Group 2 mutants (*sov*^2729^, *sov*^2^), in which the heterochromatin-promoting function of *Sov* is disrupted, exhibited a global deregulation of gene expression and derepression of transposons (FigureS6B). As expected, the *sov^ΔHBS2^* allele exhibited no impact on global gene expression. Intriguingly, telomeric transposons (HeT-A, TAHRE, TART) were upregulated, implicating the HBS2-mediated direct Sov–HP1a interaction specifically in telomere regulation (FigureS6C).

Together, the transcriptome, PEV, and complementation analyzes indicate that the transcriptional regulatory capacity of Sov is essential for the piRNA-mediated silencing, even when Sov is properly targeted to nascent transposon RNAs and is otherwise capable of promoting heterochromatin formation at the transposon locus. Conversely, while the transcriptional repression mediated by Sov alone can inhibit transcription, it is insufficient for stable transposon silencing; a Sov-dependent establishment of a heterochromatic environment at the locus is required for maintenance. This observation provides strong support for and further validates the previously proposed model, which divides transposon repression into two distinct phases: an initiation phase, mediated by RNA-dependent transcriptional repression, and a subsequent maintenance phase, driven by chromatin reorganization.^10,16^ The phenotypic analysis of the specific *sov* alleles revealed that Sov bridges these two processes, utilizing its distinct functional modules to engage in both phases of cotranscriptional transposon silencing.

## DISCUSSION

The process of TGS is executed through the coordinated action of numerous protein complexes.^49^ The recognition complex activates endogenous inhibitory processes of the cell at the transposon loci and suppresses expression at two distinct layers of gene regulation. First, these mechanisms act directly at the level of RNA metabolism to block transposon transcription. The Piwi/piRNA pathway stalls RNA polymerase II at the transposon loci, facilitating transcriptional termination and subsequent exosomal degradation of the nascent transposon RNAs.^16,19,20^ Second, the Piwi/piRNA pathway acts at the chromatin level to induce local heterochromatinization of the loci, where compact chromatin precludes further transposon transcription.^6,8,22–25^ Our results demonstrate that Sov is a central player in both silencing pathways.

Here we show that Sov functions in RNA decay at transposon loci. By binding to the SFiNX complex through Panx, Sov associates with the nascent transposon mRNA and performs RNA-based silencing that is independent of its heterochromatin-promoting activity. Specifically, Sov physically interacts with TEsup1, thereby bridging piRNA-pathway-mediated transposon RNA recognition and RNA decay. In addition, Sov contributes to transposon silencing by regulating overall heterochromatin organization. Sov promotes heterochromatinization via its specific IDRs by inducing the formation of a phase separated nuclear domain. This activity of Sov may be associated with its direct interaction with HP1a, which may facilitate the chromatin targeting of Sov activity or stabilize the phase-separated domain around specific genome regions.

### Sov regulates heterochromatin through phase separation

We demonstrate that the Sov protein directly interacts with the core heterochromatin effector HP1a, and we have identified the specific domain of Sov required for its association with heterochromatin. Multiple IDRs of Sov are capable of driving phase separation; notably, the region responsible for heterochromatic localization coincides with one of these IDRs. While IDRs in general appear to be insufficient on their own for enrichment within subnuclear domains, the IDR of Sov is nonetheless sufficient for heterochromatic accumulation through its phase-separating capacity.^50^ Phase separation is likely a prerequisite for heterochromatic localization, although we cannot exclude the involvement of other mechanisms, such as zinc-finger-mediated specific DNA binding or localization mediated by yet unknown protein-protein interactions. Although piRNA target recognition and transcription termination also involve phase separation, that of Sov is dispensable for these processes and appears to be specifically linked to heterochromatin function.

It has been shown that the Piwi–piRNA pathway triggers transposon silencing via a stepwise cascade, initiated by the loss of active histone marks and spatial repositioning of genomic regions, which acts as a precursor to the recruitment of repressive marks and the formation of a condensed chromatin state.^51,52^ However, the molecular mechanism governing where Sov integrates into this hierarchy and how it promotes heterochromatin formation remains elusive. Our previous results indicate that Sov functions downstream of chromatin-associated HP1a and stabilizes the established heterochromatin.^18^ Consistently, we have identified multiple HP1a-binding and phase-separating regions within the Sov protein. Previous studies identified the pentameric PxVxL motif as the primary sequence binding the HP1 chromo shadow domain dimer.^34^ Efficient HP1a binding, however, requires additional criteria collectively termed the HP1a Access Code (HAC).^53^ According to this molecular grammar, a canonical HAC consists of an expanded PxVxL motif within an IDR, where alternating hydrophobic residues are nested inside a basic, positively charged cluster to optimize electrostatic interactions. Although multiple HP1a binding sites (HBSs) in Sov have been confirmed *in vitro* and *in vivo*, HBS2 best satisfies the HAC. Intriguingly, deleting HBS2 alone causes no mutant phenotype, suggesting HBS redundancy in stabilizing heterochromatin. Alternatively, Sov may act independently of HP1a binding to drive phase separation through multivalent interactions, thereby establishing a molecular microenvironment that promotes heterochromatin function.

### Sov links transposon recognition to the RNA decay pathway

Our results indicate that effective transposon repression relies on the Sov-mediated recruitment of TEsup1 to the transposon transcripts via Sov-PRM/TEsup1-PRD interaction, which directs the transposon RNA toward degradation. Remarkably, the Sov interacting interface on TEsup1 coincides with the PRD that engages the PRM motifs of the nuclear exosome adaptor scaffold subunits dZfc3h1, dZcchc7 and dZcchc8.^20^ We hypothesize that the simultaneous binding of TEsup-PRD by Sov-PRM and the exosome adaptor complexes-PRMs occurs by the homo- or heterooligomerization of TEsup proteins. Although the precise stoichiometry of this complex remains uncharacterized, the PRDs of the TEsup complex are simultaneously bound by the PRMs of both Sov and the adaptor proteins to recruit the nuclear exosome to nascent transposon RNAs. In this configuration, TEsup proteins function as linkers between Sov, which specifically associates with transposon RNAs, and the NEXT, PAXT, and TRAMP adaptor complexes that target the RNA for degradation. Consequently, transposon transcripts are degraded through the SFiNX-Sov-TEsup1/2-Adaptor-nuclear exosome axis.

As an alternative hypothesis, mutually exclusive and temporally separated Sov–TEsup1 and adaptor–TEsup1 interactions could enable a binary switch-like mechanism. Such dynamic rearrangements via shared binding sites frequently regulate RNA 3’ end biogenesis and degradation.^54–57^ In this scenario, shared sites ensure that Sov initially recruits TEsup1 to nascent transposon RNAs. Upon localization, Sov releases TEsup1 to the nuclear exosome adaptor complex, which binds the target RNA via its own components to initiate degradation. This competitive switch is likely driven by differential binding affinities. Crucially, this system acts as a safety gate, ensuring TEsup1 recruits the destructive nuclear exosome machinery only after correct positioning by Sov, thereby preventing accidental host mRNA degradation.

### A model for Sov function in transposon silencing

While it is established that both the transcriptional and heterochromatic repression is indispensable for transposon silencing, a model for the coordinating these two effector processes is emerging. Sov may integrate these two pathways by acting as a molecular scaffold, through its diverse molecular activities, from which both pathways diverge. When associated with the SFiNX complex, Sov acts as an upstream element in the regulatory network of RNA metabolism. Conversely, in heterochromatin regulation, it functions as a downstream effector of chromatin-bound HP1a, serving as an integral component of a complex negative feedback loop.^18,28^ Sov is targeted to transposons via Panx binding within the SFiNX complex; however, it can localize directly to established heterochromatin at transposon loci via phase separation mediated by its heterochromatin localization domain. Previous studies have demonstrated that direct transcriptional silencing and heterochromatin deposition are temporally uncoupled, with transcriptional repression preceding heterochromatin formation.^10,16^ Consequently, Sov appears to function as a stable core component of a temporally dynamic multiprotein assembly at transposon loci, executing distinct roles throughout the silencing process.

However, many unanswered questions remain. For example, the role of the nucleolar localization of Sov and its link to transposon silencing or chromatin regulation remain unclarified. In addition, our analysis of mutant phenotypes identified distinct regions in Sov critical for transposon silencing, though their exact molecular functions remain unclear. These uncharacterized regions might participate in linking Sov to pathways such as SUMOylation-dependent transcription termination or DNA-based silencing mechanisms.

In summary, our results suggest that mediating the connection between the SFiNX complex and the heterochromatin machinery is not the sole function of RNA-tethered Sov in the transposon repression. In line with previous hypotheses, when recruited by Panx to transposon loci, Sov does not primarily act at the chromatin level to repress transcription, but instead engages a transcriptional repressor mechanism that acts directly on the transcription process.^16^ Subsequently—or in parallel—heterochromatinization induced by other regions of Sov contributes to the maintenance of repression and the establishment of stable transposon silencing. Consequently, Sov represents a unique element within the cotranscriptional transposon silencing pathway, exerting a direct role at both the RNA-targeted and chromatin-level regulatory layers.

## MATERIALS AND METHODS

### Plasmid construction

*In vitro* synthesized *sov* CDS (GeneUniversal, Newark, DE) or FlyFos018439 fosmid was used as a template to amplify the required Sov fragments.^58^ The amplified fragments were cloned into linearized vectors using gene assembly (New England Biolabs, E5520). For TagEGFP and mCherry tagging of Sov, PCR-amplified TagEGFP and mCherry were inserted into the NotI/BamHI sites of the pUASP-K10-attB plasmid.^59^ For N-terminal tagging, PCR-amplified *sov* fragments carrying an SV40 NLS were cloned into BamHI-digested pUASP-K10-attB-EGFP and pUASP-K10-attB-mCherry plasmids. For C-terminal λN tagging, the pUASP-K10-attB-λN-NLS plasmid was first generated by inserting the *in vitro* synthesized λN-NLS fragment into the NotI/BamHI sites of the pUASP-K10-attB vector. Then, *sov* fragments were cloned into the NotI site of the pUASP-K10-attB-λN-NLS vector. For C-terminal tagging of Sov fragments with mCherry:Cry2, the pHR-mCh:Cry2olig (Addgene101222) plasmid was PCR-amplified in two parts, and the two plasmid fragments were assembled together with PCR-amplified *sov* fragments. For GST tagging, PCR-amplified *sov* fragments were cloned into BamHI/XhoI-digested pGEX-6P-1 plasmid. Various mutant *sov* fragments were generated by *in vitro* mutagenesis (Agilent Technologies, 210518). For His-tagged HP1a production, the HP1a coding region was PCR-amplified from LD10408 and inserted into the NdeI/BamHI sites of pET16b. For production of His-tagged Sov fragments, Sov(819-1299) and Sov(157-816) coding sequences were PCR-amplified and inserted into the NdeI/BamHI sites of pET16b. For Y2H screening, LD10408, LD18231, SD08161, FI04411, LD36051 cDNA clones, and FlyFos018439 fosmid were used to amplify coding sequences of HP1a, Panx, Nxf2, Piwi, TEsup1, and Sov, respectively. Coding sequences of TEsup2 were amplified from ovarian total cDNA. PCR fragments were inserted into pGAD424 (BglII site) and pGBT9 (SalI site). All constructs were sequence verified (Eurofins Genomics, Ebersberg, Germany).

### Localization and microscopy

For live imaging experiments, ovaries from 1-day-old *adult* females were dissected in PBS and separated into individual ovarioles. Nurse cell nuclei of stage 7–9 egg chambers were analyzed in glass-bottom Petri dishes (Ibidi, 81218-200) containing Schneider’s medium supplemented with fetal bovine serum to 10%. Confocal imaging was performed using a Zeiss LSM800 confocal microscope. To assess the subnuclear localization of mCherry-or TagEGFP-tagged Sov fragments, nurse cell nuclei from stage 7–8 egg chambers were imaged in Airyscan SR mode using a Zeiss Plan-Apochromat 63×/1.4 NA oil immersion objective. For co-localization analysis, fluorescent intensity profiles were measured with ImageJ. For tethering assays, stage 5-6 egg chambers were imaged using a Zeiss Plan-Apochromat 20×/ 0.8NA objective.

### Thermostability assay

Purified His-tagged Sov(819–1299) and Sov(157–816) protein fragments were diluted in PBS to a final concentration of 1 mg/ml and were subjected to 95°C for temperature for 15 minutes followed by centrifugation at 15000g that separated the insoluble fraction, representing the thermo-sensitive population, from the soluble proteins, representing the thermo-resistant fraction. Purified EGFP protein was used as control.

### optoDroplet assay

HEK293T cells (ATCC, CRL-3216) were maintained in Dulbecco’s modified Eagle’s medium (Capricorn Scientific, DMEM-HPA) supplemented with 10% fetal bovine serum (Capricorn Scientific, 10-FBS-HI-12F) and 1% penicillin-streptomycin (Capricorn Scientific, PS-B). Cells were cultured at 37°C in a humidified incubator with 5% CO_2_. For live-cell imaging, 10^4^ cells were plated onto 8-well glass-bottom µ-Slides (Ibidi, 80827) 24 hours prior to transfection. optoDroplet plasmids were transfected using jetOptimus reagent (Sartorius, 101000051). Immediately before imaging, the culture medium was replaced with Live Cell Imaging Solution (Invitrogen, A14291DJ). Images were acquired using a VisiScope spinning disk confocal microscope equipped with a Yokogawa CSU-W1 confocal scanning unit and Andor Zyla 4.2 PLUS sCMOS cameras (Visitron Systems). For the imaging, a PlanApo N 60x Oil 1.42NA Objective (Olympus) was used. Optogenetic activation was induced via 488 nm laser pulses for 1s. Area covered by the droplets was measured with ImageJ (TableS1).

### Protein production and purification

For production of the His-tagged HP1a, Sov(819–1299) and Sov(157–816) proteins, the pET16b plasmid constructs were transformed in E. coli strain BL21(DE3) SixPack and induced by 0.5 mM IPTG at 37 °C for 4 hours.^60^ Cells were lysed in lysis buffer [100 mm NaH_2_PO_4_ × H_2_O pH 8, 150 mM NaCl, 5 mM Imidazol, 7× EDTA-free protease inhibitor cocktail (Roche, 04693132001)] a Covaris M220 focused ultrasonicator (Covaris settings: peak power: 75W, duty factor 10%, duration: 420s, cycles/burst:25). Triton X-100 was added to the lysate to a final concentration of 1%, followed by incubation on ice for 30 min. Talon Metal Affinity Resin (Takara, 635502) beads were equilibrated using five bead volumes of lysis buffer and was incubated with lysate for 2 h at 4 °C. After three washes with washing buffer (100 mM NaH_2_PO_4_ × H_2_O pH 8, 300 mM NaCl, 20 mm Imidazol), His-tagged proteins were eluted with elution buffer (100 mM NaH_2_PO_4_ × H_2_O pH 8, 300 mM NaCl, 300 mM Imidazol). Elution buffer was exchanged for storing buffer (50 mM Tris–HCl, pH 7.4, 150 mM NaCl, 1 mM DTT, 5% glycerol) using a PD-midiTrap G-25 column (GE Healthcare, 28918008). The protein solution was concentrated using a 3 kDa MWCO Amicon Ultra-4 Centrifugal Filter (Millipore, UFC8003) and stored at −80 °C.

### GST pull-down assay

For production of GST or GST-tagged proteins, pGEX-6P-1 constructs were transformed in E. coli strain BL21 and induced by 0.5 mM IPTG at 18 °C O/N. Cells were lysed in lysis buffer (50 mM Tris–HCl, pH 7.4, 5 mM DTT, 50 mM NaCl, 5 mM EDTA, 10% glycerol, 25× protease inhibitor cocktail (Roche, 11697498001) using a Covaris M220 focused ultrasonicator. Triton X-100 was added to the lysate to a final concentration of 1%, followed by incubation on ice for 30 min. Glutathione Sepharose 4B (GE Healthcare) beads were equilibrated using five bead volumes of lysis buffer and were incubated with lysates for 2 h at 4 °C. After three washes with washing buffer I (50 mM Tris–HCl, pH 7.4, 5 mM DTT, 400 mM NaCl, 10% glycerol), BSA and purified His-HP1a were added to a final concentration of 500µg/ml and 1µg/ml, respectively, and the mixture was incubated for 2 h at 4 °C. After washing three times with the washing buffer II (50 mM Tris–HCl, pH 7.4, 5 mM DTT, 50 mM NaCl, 5 mM MgCl_2_, 10 mM KCl, 5% glycerol) the proteins were eluted by incubating the beads with 2× SDS/PAGE buffer for 5 min at 95 °C. The GST-tagged bait proteins were visualized with Coomassie blue staining, the His-HP1a protein was detected by western blot.

### Western blot

Proteins were separated by SDS-polyacrylamide gel electrophoresis and subsequently transferred onto Immobilon-P PVDF membranes (Millipore, IPVH00010). Membranes were incubated in mouse anti-HP1a (1:2000, C1A9, DSHB) primary antibodies at 4 °C overnight. Horseradish peroxidase (HRP) -conjugated anti-mouse IgG (1:5000, Sigma, A2304) was applied as a secondary antibody for 1h at room temperature. Immunocomplexes were visualized with Immobilon Western Chemiluminescent HRP Substrate (Millipore, WBKLS0100) using the UVITEC Alliance Q9 system.

### Yeast Two-Hybrid Assay

Yeast Two-Hybrid assays were performed using the Matchmaker two-hybrid system (Clontech, K1605).^61^ The binding (pGBT9) and activation (pGAD424) domain fusion constructs were co-transformed into the *Saccharomyces cerevisiae* reporter strain PJ69-4A. Transformants were initially selected on synthetic complete double drop-out medium lacking leucine and tryptophan (SC-Leu/-Trp, Merck, Y0750). To assess protein–protein interactions, single colonies were resuspended in sterile water at equal densities and spotted onto SC-Leu/-Trp plates to confirm viability, and onto triple drop-out plates lacking leucine, tryptophan, and histidine (SC-Leu/-Trp/-His,Merck, Y2146) to monitor HIS3 gene expression. All plates were incubated at 30 °C for 3–5 days before scoring.

### RNA purification, real-time quantitative PCR (RT-qPCR), mRNA sequencing, and computational analysis

For RT-qPCR and RNA-seq, total RNA was isolated from 15 pairs of ovaries of 2-day-old adult females in three biological replicates using the Reliaprep RNA Tissue Miniprep System (Promega, Z6111). For RT-qPCR, 1 µg total RNA was reverse-transcribed with random primers using the First Strand cDNA Synthesis kit (Thermo Scientific, K1612). Three technical replicates were used for qPCR with Maxima SYBR Green/ROX qPCR Kit (Thermo Scientific, K0221) on a Rotor-Gene Q System (Qiagen). Gene expression was quantified using the comparative C_T_ method (ΔΔC_T_ method). Primers used in PCR reactions are listed in TableS2. RNA-seq was performed by Novogene (Munich, Germany). RNA-seq libraries were constructed following polyA enrichment and sequencing was performed on the Illumina NovaSeq X Plus system. Approximately 20 million reads per sample (150-bp, paired-end reads) were obtained. Raw reads were processed using Trimmomatic-0.39, and then aligned to the reference sequence using Kallisto-0.48.0.^62,63^ Since sequences of transposable elements were not found in dmel-all-transcript-r6.55.fasta in FlyBase, a list of the consensus transposon sequences in were obtained from (https://github.com/bergmanlab/drosophila-transposons). and used as reference data. Gene-level count matrices were created for use with DESeq2 by importing the quantification data using tximport^64^. For differential analyzes the DESeq2-1.42.1 R package was used.^65^ All raw sequence data have been deposited in the SRA (www.ncbi.nlm.nih.gov/sra) under accession number PRJNA1492023.

### Chromatin Immunoprecipitation-qPCR

For each sample, ovaries were dissected from 30 females in ice-cold PBS and fixed in PBS+1% paraformaldehyde for 10 min at room temperature. Crosslinking was quenched by the addition of glycine to a final concentration of 125 mM for 5 min at room temperature. Ovaries were subsequently washed twice in PBS for 5 min at room temperature and collected by centrifugation at 3,000 rpm for 1 min at 4 °C. Chromatin isolation, shearing, and immunoprecipitation were performed using the qPCR & Sequencing Chromatrap ChIP Kit (Porvair Sciences; 500274) according to the manufacturer’s instructions with minor modifications. Briefly, ovaries were resuspended in Hypotonic Buffer and incubated at 4 °C for 10 min. Following centrifugation at 3,000 rpm for 1 min at 4 °C, samples were resuspended in Lysis Buffer and homogenized using a plastic pestle, followed by incubation on ice for 10 min. Chromatin was sheared using a Covaris M220 Focused-ultrasonicator at 4 °C to obtain DNA fragments ranging from 100–500 bp (Covaris settings: peak power: 75W, duty factor 10%, duration: 1500s, cycles/burst:200). Sonicated lysates were cleared by centrifugation at 14,000 rpm for 15 min at 4 °C, and the supernatant containing sheared chromatin was collected. Sheared chromatin was diluted with Column Conditioning Buffer, and 4 µg chromatin was incubated with 5 µg of mouse anti-Histone H3 tri-methyl K9 (H3K9me3) antibody for 4 h at 4 °C (Porvair Sciences; 700003). Immunoprecipitated chromatin was subsequently loaded onto Protein G Chromatrap spin columns, and washing and elution steps were performed according to the manufacturer’s instructions. Reverse crosslinking was carried out overnight at 65 °C in 100 mM NaHCO_3_ and 200 mM NaCl. Samples were then treated with RnaseA (2 mg/ml) at 37 °C for 30 min, followed by ProteinaseK (2 mg/ml) treatment at 37 °C for 60 min. DNA was purified using the PCR Cleanup Kit according to the manufacturer’s instructions (Geneaid, DFC100). The quantity of isolated DNA was determined using the Qubit 1X dsDNA HS Assay Kit (Thermo Scientific, Q33230) on a Qubit 4 fluorometer. Subsequently, qPCR was performed using the Maxima SYBR Green/ROX qPCR Kit (Thermo Scientific, K0221) on a Rotor-Gene Q System (Qiagen). Fold enrichment was calculated using the ΔΔC_T_ method according to.^22^ Primers used in PCR reactions are listed in TableS2.

### Proximity labelling

#### Sample preparation

Proximity labelling was performed as described previously.^66^ In brief, females simultaneously expressing *eggless* regulator-driven TurboID biotin ligase fused to nuclear GFP-nanobody (TurboID-vhhGFP-3xHA-NLS) and Ubi>NLS:EGFP (control-TurboID) or *nos*-Gal4-driven Sov(337–521):EGFP (Sov-TurboID) were used. Three Sov-TurboID and four control-TurboID samples were prepared. Prior to tissue collection, flies were fed for 16 h with yeast paste supplemented with biotin (Sigma, B4501) to 100 µM. Ovaries were dissected from approximately 60 flies (corresponding to ∼100 µl tissue volume) and stored at −80 °C. Dissected ovaries were washed once with ice-cold PBS and homogenized in 1 ml pre-extraction buffer (10 mM Tris-HCl pH 7.5, 2 mM MgCl_2_, 3 mM CaCl_2_, 0.5% NP40, cOmplete Protease Inhibitor Cocktail (Roche, 11697498001), 10% glycerol) using a Dounce homogenizer, followed by incubation at 4 °C with nutation for 15 min. Samples were centrifuged at 20000g for 5 min at 4 °C to remove cytoplasmic contaminants and enrich for nuclei. The nuclear-enriched pellet was resuspended in 100 µL lysis buffer (50 mM Tris-HCl pH 7.5, 150 mM NaCl, 0.1% SDS, 0.5% Na-Deoxycholate, 1% Triton-X, 1 mM DTT, Benzonase, cOmplete Protease Inhibitor Cocktail) and further homogenized using an electric plastic pestle for 20 s on ice. Subsequently, 900 µl lysis buffer was added, and samples were additionally homogenized by further douncing (20 times). Lysates were incubated at 4 °C with nutation for 1–2 h and cleared twice by centrifugation at 18000g for 10 min each. For affinity purification, 80 µl Pierce magnetic streptavidin beads (Thermo Fisher Scientific, 88817) were pre-equilibrated in 1 ml lysis buffer. Cleared lysates were incubated with pre-equilibrated streptavidin beads for 1 h at 4 °C. Beads were washed sequentially with lysis buffer for 10 min at 4 °C, lysis buffer containing 2% SDS for 10 min at room temperature, wash buffer I (50mM HEPES pH 7.5, 500 mM NaCl, 1 mM EDTA, 0.1% Na-Deoxycholate, 1% Triton-X) for at least 10 min, and wash buffer II (10mM Tris-HCl pH 7.5, 250 mM LiCl, 1 mM EDTA, 0.5% Na-Deoxycholate, 1% NP40) for at least 10 min, followed by five washes in TBS (20 mM Tris-HCl pH 7.5, 137 mM NaCl).

#### Mass spectrometry analysis

Samples were processed using Gel-aided Sample Preparation (GASP) as previously described^67^. Briefly, protein samples were immobilized in an acrylamide gel matrix, followed by in-gel reduction, alkylation, and tryptic digestion. Peptides were extracted from the gel using aqueous acetonitrile containing formic acid, pooled, dried, and reconstituted prior to LC– MS/MS analysis. Peptide samples were loaded onto Evotips and analyzed using an Evosep One liquid chromatography system employing the 88 min gradient corresponding to the 15 samples/day method. Eluting peptides were analyzed on an Orbitrap Fusion Lumos mass spectrometer equipped with a FAIMS Pro interface applying multiple compensation voltages. MS1 spectra were acquired in the Orbitrap at high resolution, while MS/MS spectra were generated by higher-energy collisional dissociation (HCD) and detected in the ion trap. Raw data were processed using Proteome Discoverer (version 3.0) with the Byonic search engine. Spectra were searched against the UniProt Swiss-Prot *Drosophila melanogaster* database supplemented with a common contaminant database. Trypsin specificity was applied allowing up to two missed cleavages. Precursor and fragment mass tolerances were set to 5 ppm and

0.6 Da, respectively. Carbamidomethylation of cysteine was specified as a fixed modification. Variable modifications included protein N-terminal acetylation, methionine oxidation, N-terminal methionine loss, and N-terminal pyro-glutamate formation. A limited number of variable modifications per peptide was allowed. Label-free quantification was performed using precursor ion intensities. Protein abundances were calculated from summed peptide abundances using the top three peptides per protein (Top N = 3).

#### Mass spectrometry data analysis

Protein abundance data were analyzed using the Perseus computational platform (v.2.1.3.0).^68^ The dataset was cleaned by filtering out non-*Drosophila* contaminants. Only proteins detected across all biological replicates in either the control or the Sov-TurboID groups were retained for downstream analysis. Missing values were imputed using the Quantile Regression Imputation of Left-Censored data (QRILC) method. Differential expression analysis was performed using edgeR. Log_2_-transformed intensity values were normalized via the Trimmed Mean of M-values (TMM) method, and statistical significance was evaluated using the Likelihood ratio test (TableS3).

### AlphaFold3 screening

To identify Sov(337–521) interactors, pools of candidate partner proteins were selected from the following sources: (1) significantly enriched Sov(1–1500) interactors identified by co-immunoprecipitation^16^, (2) significantly enriched Sov(337–521) interactors determined via TurboID (this study), (3) proteins involved in RNA metabolic processes required for transposon silencing^19,20^, and (4) the structured domains of these proteins (TableS4). AlphaFold structures of the analyzed full length candidate proteins were retrieved from AlphaFoldDB using the AlphaFoldDB Structure Extractor.^69,70^ From these protein structures confidently predicted structured regions (pLDDT scores between 70 and 100) spanning more than 100 amino acids were extracted. Adjacent structured regions located closer than 100 amino acids to each other were merged. The sequences of both full-length proteins and their structured protein fragments were retrieved from the UniProt database and were pairwise tested with Sov(337–521) using the AlphaFold3 server (TableS4). Calculations were performed with default settings.^71^ For each prediction, five structural models were generated and quantified. For each model, pDockQ2 values were determined using the af_analysis software package.^72,73^

### Drosophila strains

*Drosophila* stocks were maintained at 25 °C on standard medium. Transgenes were integrated into the attP40 landing site (BDSC #36304) by standard methods using the PhiC31 integrase-mediated site-specific transgenesis (BDSC #40161). Transgenes were driven by *nos*-Gal4:VP16 (BDSC #4937), α*Tub67C*-Gal4:VP16 (mat-Tub-Gal4, BDSC #7063), or α*Tub67C*-GAL4:VP16, *osk*-GAL4:VP16 (TOsk-Gal4, VDRC#314033) drivers. In the transcriptional reporter assay, the mTub>EGFP_5xBoxB_SV40 [attP2] (VDRC #313408) strain was used to test the silencing capacity of λN-tagged Sov fragments.^22^ Fib:RFP stock was obtained from Shelby Blythe.^74^ shRNA strains used in desilencing experiments are listed in TableS8. Second-chromosome shRNA lines were expressed using the mat-Tub-Gal4 driver, whereas third-chromosome lines were driven by TOsk-Gal4.^75^ For the analysis of *sov* mutant phenotypes *sov^ΔHBS2^*, *sov*^1216^, *sov*^1929^, *sov*^2627^, *sov*^3032^ (this study), the *sov*^2^ (Bl#4611) and *sov^def^*^1^ alleles were used.^18^

### Generation of *sov* mutant *Drosophila* strains

Mutant *Drosophila* strains were generated by the CrispR/Cas9 method. The gRNA target sites were selected using the flyCRISPR Optimal Target Finder tool.^76^ To generate the *sov^ΔHBS2^* allele, oligonucleotides corresponding to the gRNA target site sequence (SST1) were annealed, phosphorylated with T4 Polynucleotide Kinase (NEB), and ligated into BbsI-digested pCFD5 vector. A single-stranded oligonucleotide donor (ssdonor1) was used as repair template to introduce the 15 bp deletion into the *sov* locus (TableS5). The pCFD5-SST1 plasmid (250 ng/μl) and the ssdonor1 repair template (500 ng/μL) were co-injected into vas>Cas9 embryos. The progeny of injected flies were used to establish 250 balanced stocks, which were screened by PCR for heteroduplexes in the region flanking HBS2.^77^ Proper deletion of the HBS2 coding sequence was confirmed by sequencing. To generate *sov*^1216^, *sov*^1929^, *sov*^2627^, and *sov*^3032^ deletion alleles, pairs of gRNA target site sequences (SSTs) (TableS5) were PCR-amplified and cloned into BbsI-digested pCFD5 plasmid using Gibson Assembly (NEB, E5510).^76^ Transgenic flies were generated by integrating the pCFD5-SST plasmids expressing the appropriate gRNA pairs into the attP40 docking site. *Sov* deletions were generated in *w*^1118^/Y; Vas>Cas9 / attP40-SST males. Balanced stocks carrying mutagenized X chromosomes were established and screened for deletions by PCR. In-frame deletions were confirmed by sequencing.

### Complementation analysis, fertility test, PEV assay

For complementation analysis, females carrying different *sov* mutant alleles were crossed with OregonR males. Egg-laying assays were performed using groups of five females, and the numbers of deposited and hatched eggs were quantified daily over five consecutive days. A minimum of three independent biological replicates was analyzed for each allelic combination. Based on the number of eggs laid and hatched, the phenotypic outcomes of each pairwise *sov* allelic combination were assigned to one of six categories of increasing severity (TableS6). Correlations among *sov* alleles were assessed by Kendall’s τ correlation analysis in R using the phenotype category data. Correlation coefficients were visualized as a heatmap generated with the pheatmap R package, and hierarchical clustering was performed using the Ward.D2 method. For the position-effect variegation (PEV) assay, *sov* mutant alleles were combined with the *w^m^*^4h^ allele, and 4-day-old adult females were analyzed. Eyes were imaged using a Leica MZ FLIII stereomicroscope, and image stacks were processed using the CombineZP software. To examine ovarian morphological phenotypes, ovaries were dissected from 3-day-old adult females, fixed in 4% formaldehyde, and stained with DAPI (1:500; Sigma, D9542). Imaging was performed using a Zeiss LSM800 confocal microscope equipped with a Plan-Apochromat 20×/0.8 NA objective.

## Supporting information

The area covered by the ligth induced condensates in the optoDroplet assay

Primer sequences used for RT-qPCR and ChIP-qPCR

Mass spectrometry data of the Sov(337-521)-TurboID proximitome

Confidence metrics from the pairwise interaction screen between the Sov(337-521) fragment and selected candidate proteins or protein fragments

Repair template and gRNA target site sequences used to generate Drosophila sov alleles

Female fertility tests and phenotype categories of sov mutant allelic combinations

Normalized RNA-seq read counts generated by DESeq2 from sov mutant ovaries

Drosophila stock used in this study

## SUPPLEMENTAL FIGURE LEGENDS

**Figure S1.**
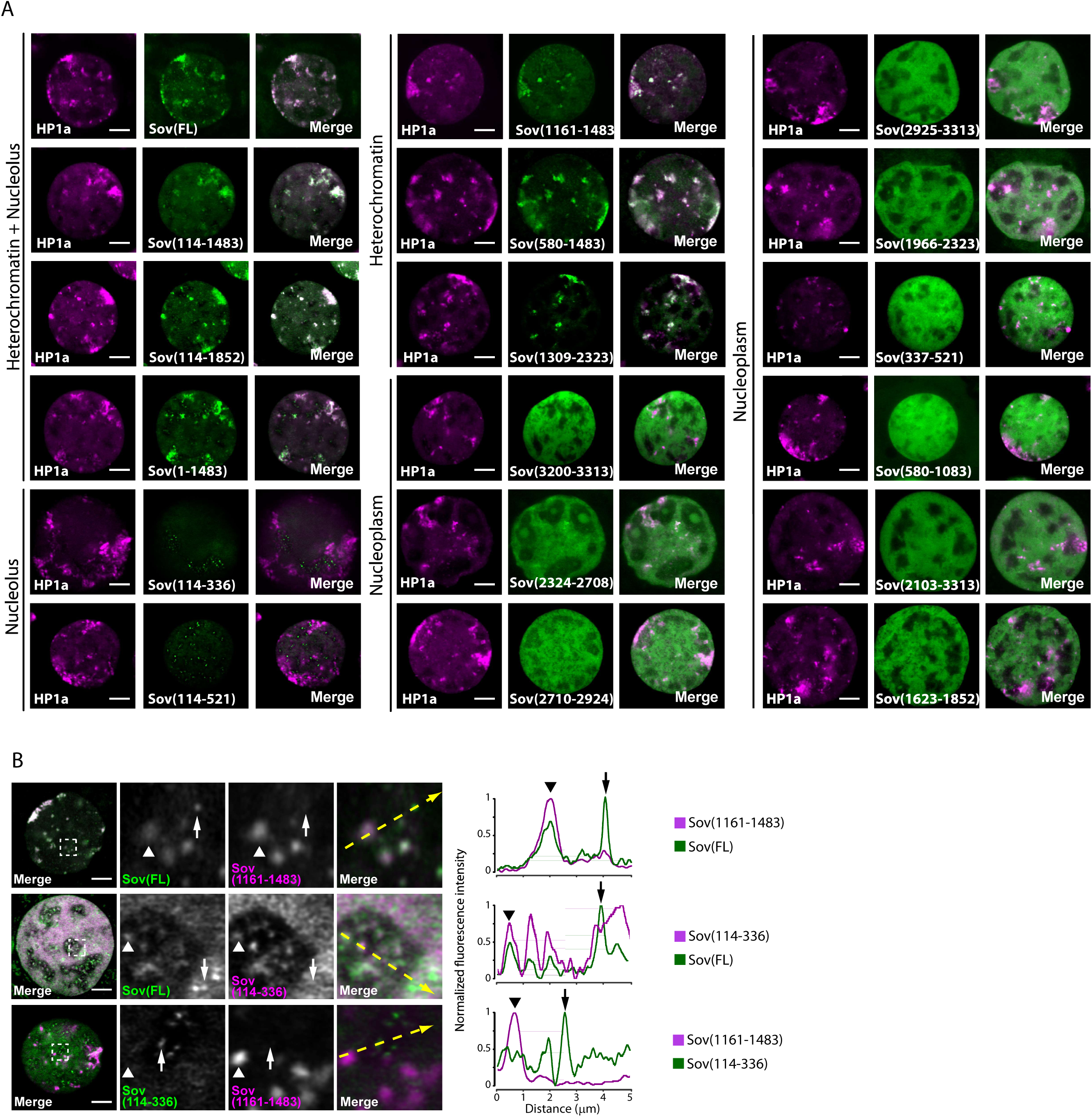
Subnuclear localization pattern of various Sov protein fragments, Related to Figure 1. **(A)** Colocalization of EGFP-tagged FL-Sov and Sov fragments with HP1a:RFP. Confocal microscopy images display germ cell nuclei. Scale bars represent 5 µm. **(B)** Confocal images display germ cell nuclei testing the colocalization of FL-Sov:EGFP with Sov(114–336):mCherry or Sov(1161–1483):mCherry, and Sov(114–336):EGFP with Sov(1161–1483):mCherry. Confocal microscopy images display germ cell nuclei; higher-magnification views of the dashed boxes are shown. Right panels present fluorescence intensity profiles quantified along yellow dashed arrows. Arrowheads and arrows mark corresponding positions between the micrographs and the intensity profile diagrams. Scale bars represent 5 µm.

**Figure S2.**
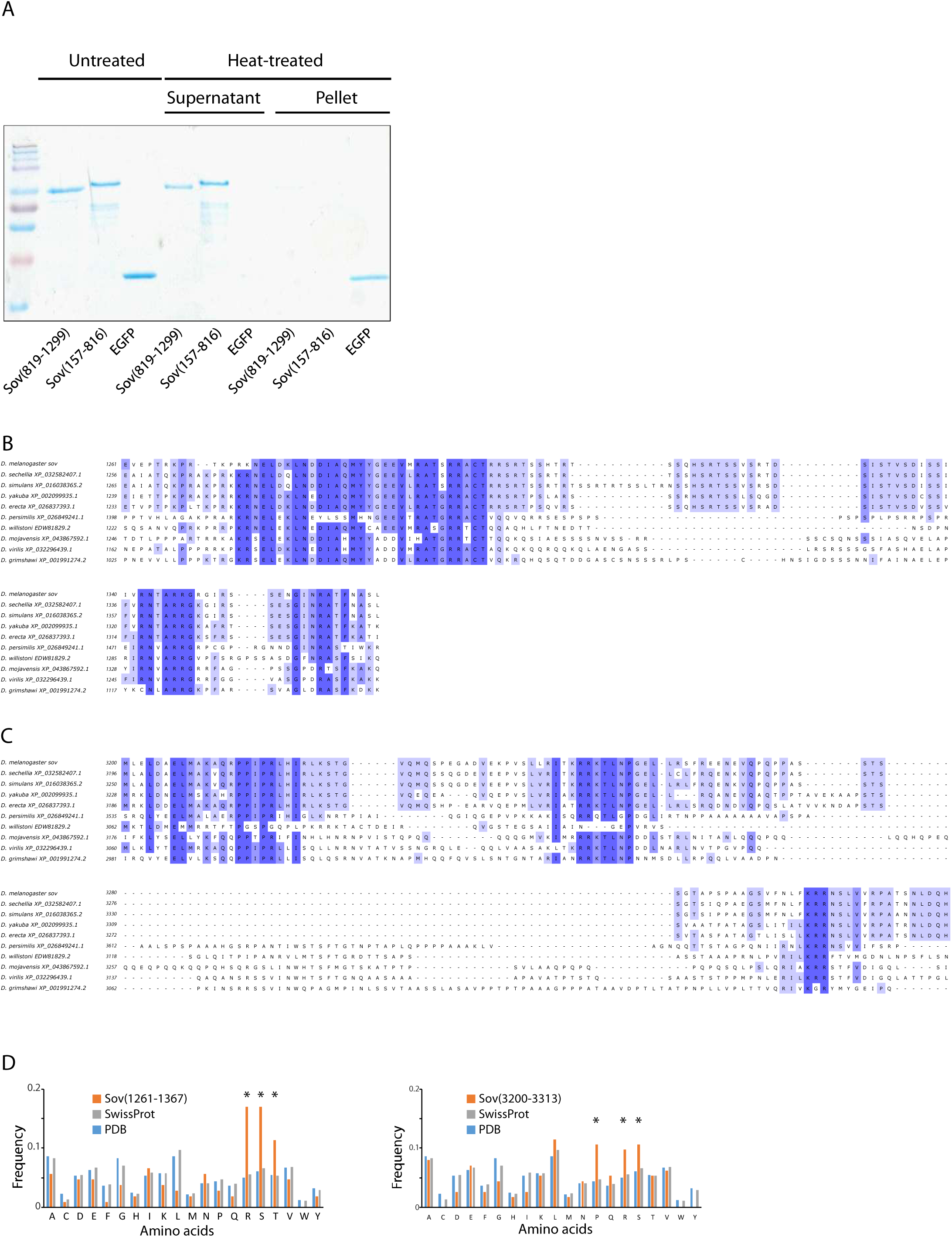
Evolutionary, compositional, and biochemical characterization of Sov disordered regions, Related to Figure 2. **(A)** Thermostability assay of disordered Sov fragments. Coomassie-stained gel showing the distribution of purified Sov(819–1299) and Sov(157–816) fragments. Following sedimentation, heat-treated disordered Sov fragments remain soluble in the supernatant, whereas the structured EGFP control precipitates into the pellet. **(B,C)** Multiple sequence alignments showing the evolutionary conservation of Sov(1261– 1367) (B) and Sov(3200–3313) (C) across representative *Drosophila* species. Shading intensity corresponds to the degree of amino acid conservation. **(D)** Amino acid frequency analysis of the Sov(1261–1367) (left) and Sov(3200–3313) (right) fragments compared to the baseline distributions in the SwissProt (grey bars) and PDB (blue bars) databases.^79^ Asterisks indicate enriched amino acid residues.

**Figure S3.**
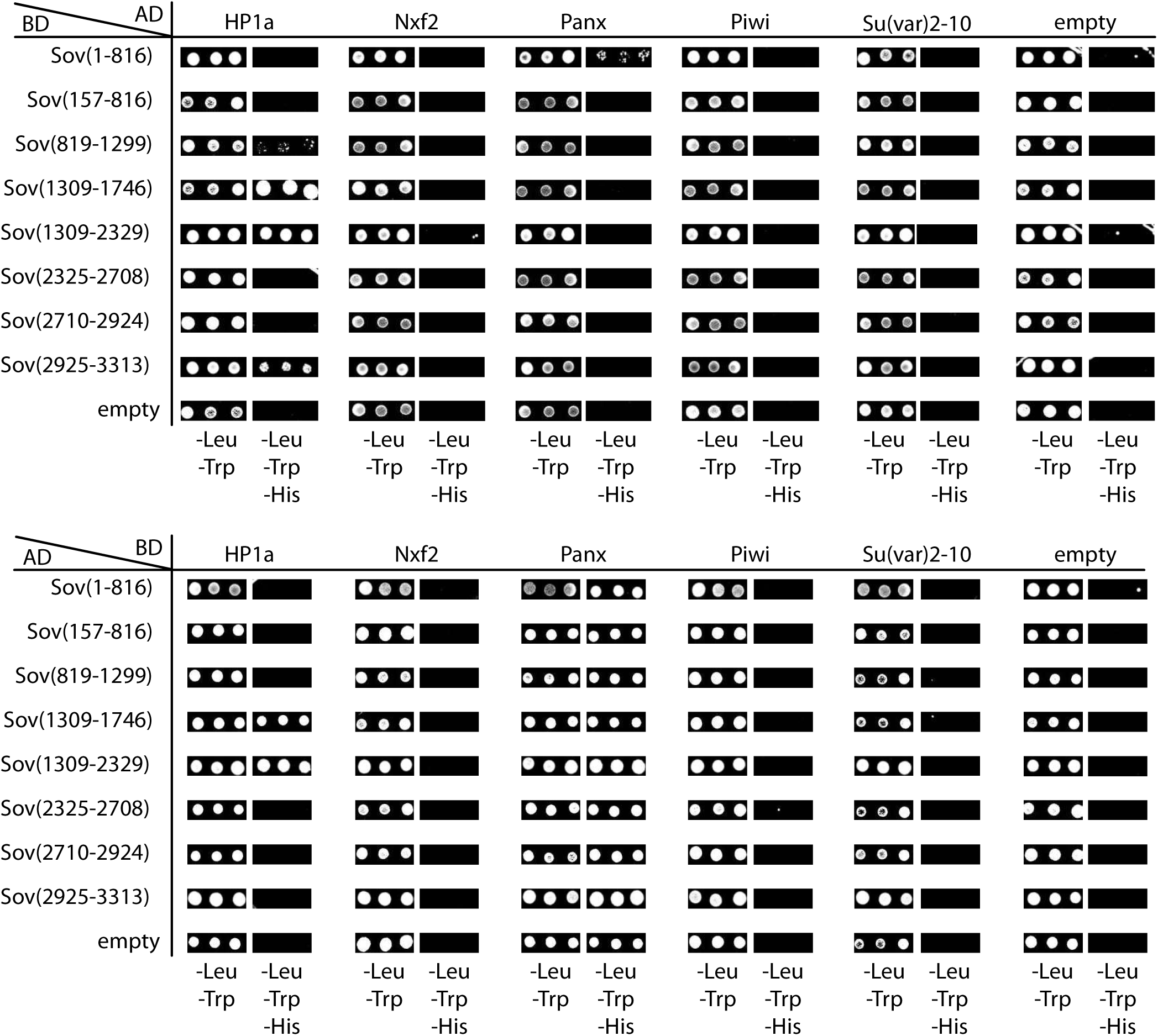
Mapping of Sov interactions with cotranscriptional silencing factors by Y2H assay, Related to Figure 3. Y2H analysis testing direct physical interactions between the indicated fragments of Sov and key heterochromatin/piRNA pathway factors. Independent panels represent a swap of the fusion configurations to rule out vector-specific artifacts. Data from three independent replicates are shown. Plasmids lacking inserts (empty) served as negative controls to monitor for background auto-activation.

**Figure S4.**
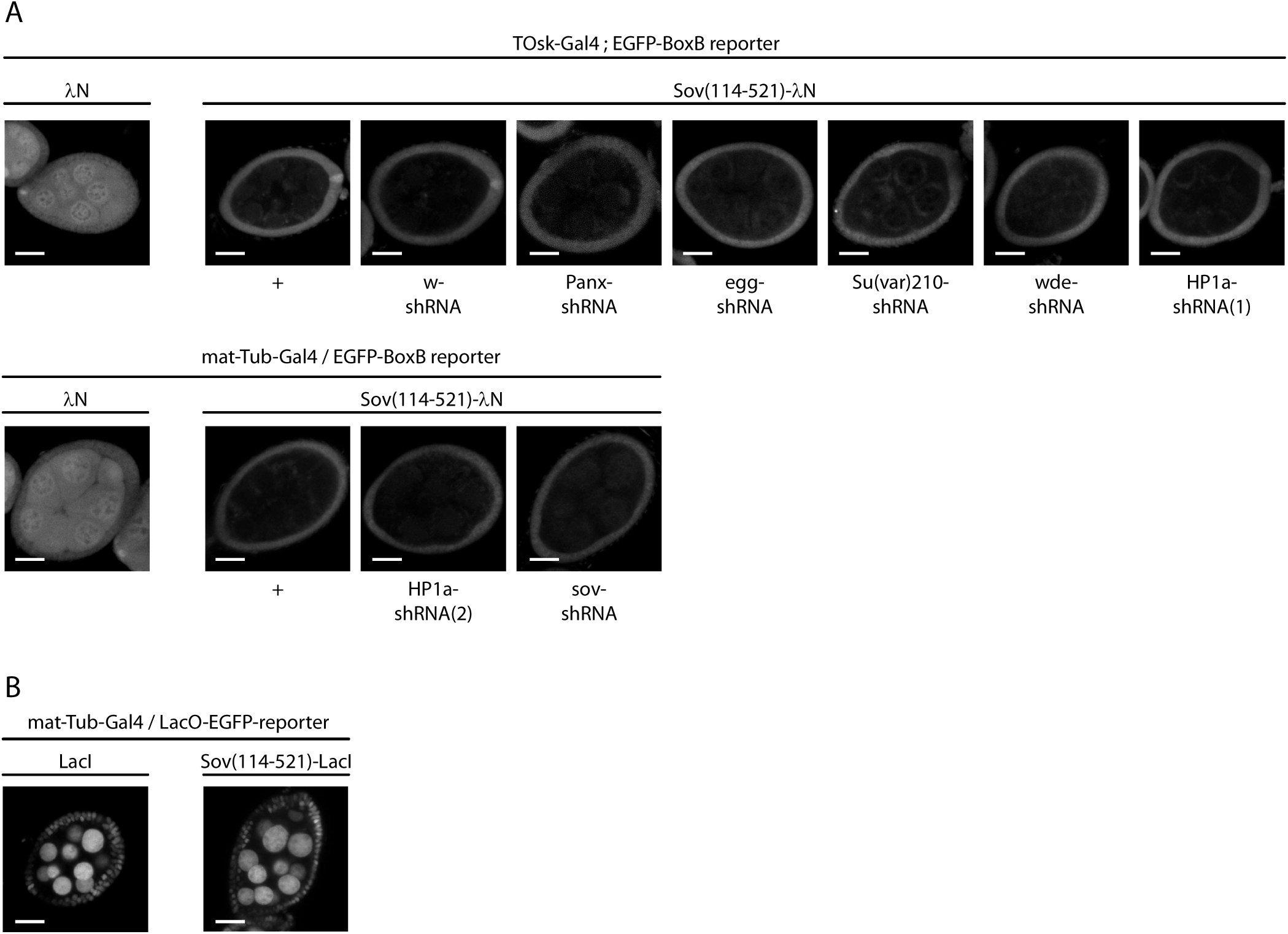
Sov-mediated tethering assays in Drosophila ovaries, Related to Figure 4. **(A)** Confocal microscopy images showing EGFP-BoxB reporter fluorescence in egg chambers. Expression of the λN-tagged Sov(114–521) fragment driven by the germline-specific mat-Tub-Gal4 or the TOsk-Gal4 drivers results in reduced EGFP reporter expression, which is not modified by the simultaneous silencing of the indicated heterochromatin regulators via shRNA constructs. Expression of the *w*-shRNA served as a negative control. Scale bars represent 20 µm. **(B)** Silencing assay based on DNA-tethering. Confocal microscopy images showing GFP fluorescence of the constitutively expressed LacO-GFP:NLS reporter.^22^ Expression of the LacI-tagged Sov(114–521) fragment driven by the germline-specific mat-Tub-Gal4 driver has no effect on GFP reporter expression. Expression of the LacI-tag served as a negative control. Scale bars represent 20 µm.

**Figure S5.**
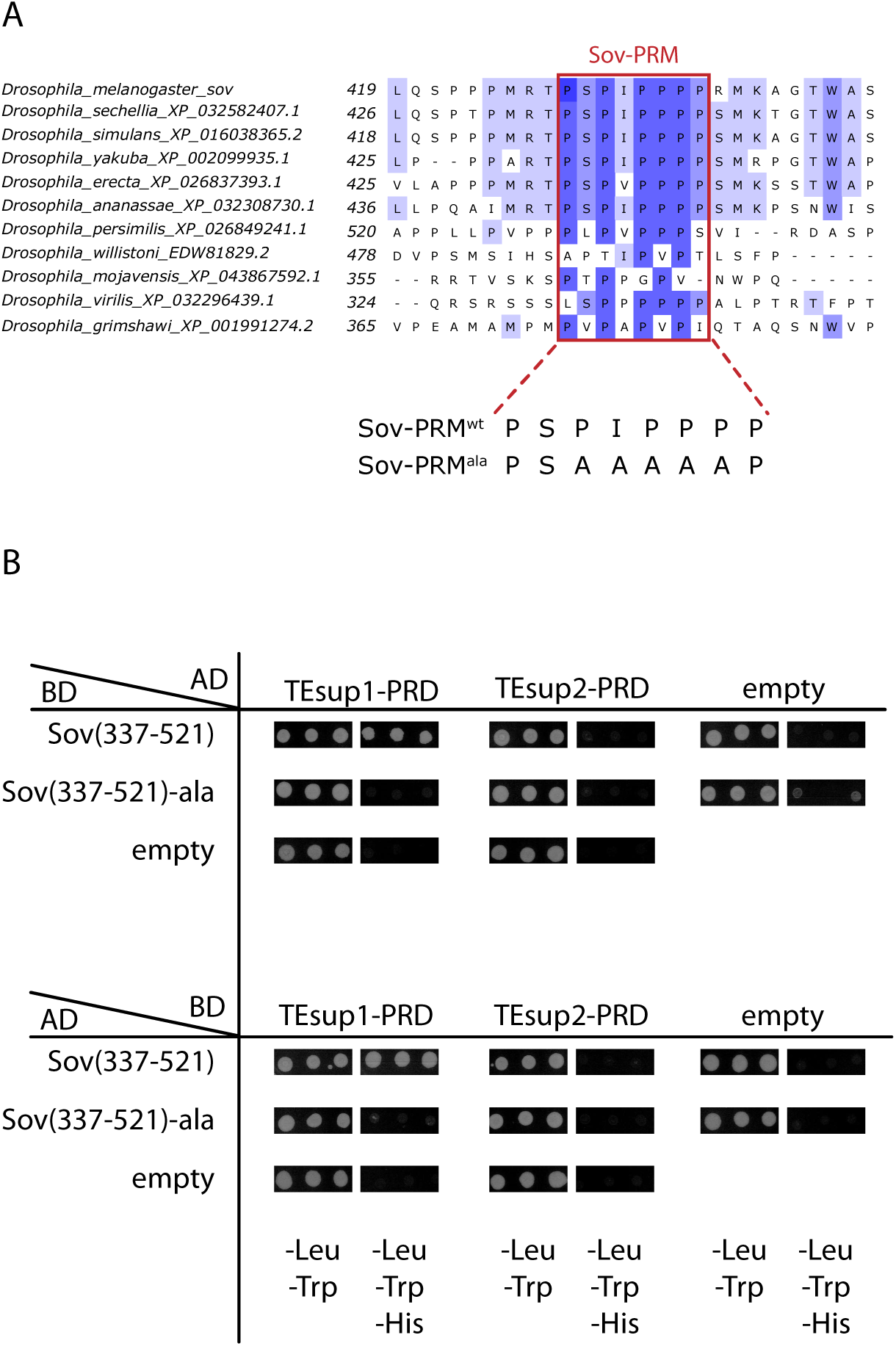
Y2H confirms Sov-PRM binding to TEsup1-PRD, Related to Figure 5. **(A)** Multiple sequence alignment of the Sov proline-rich motif (Sov-PRM) across indicated *Drosophila* species. The core consensus sequence (residues 431–436 in *D. melanogaster*) is highlighted. The amino acid substitutions to generate Sov(337-521)ala, used for the Y2H screen, are shown below. **(B)** Y2H analysis testing direct physical interactions of Sov(337–521) and Sov(337– 521)ala with TEsup1 and TEsup2. For each test, colonies from three independent replicates are shown. Independent panels represent a swap of the fusion configurations to rule out vector-specific artifacts. Plasmids lacking inserts (empty) served as negative controls to monitor for background auto-activation.

**Figure S6.**
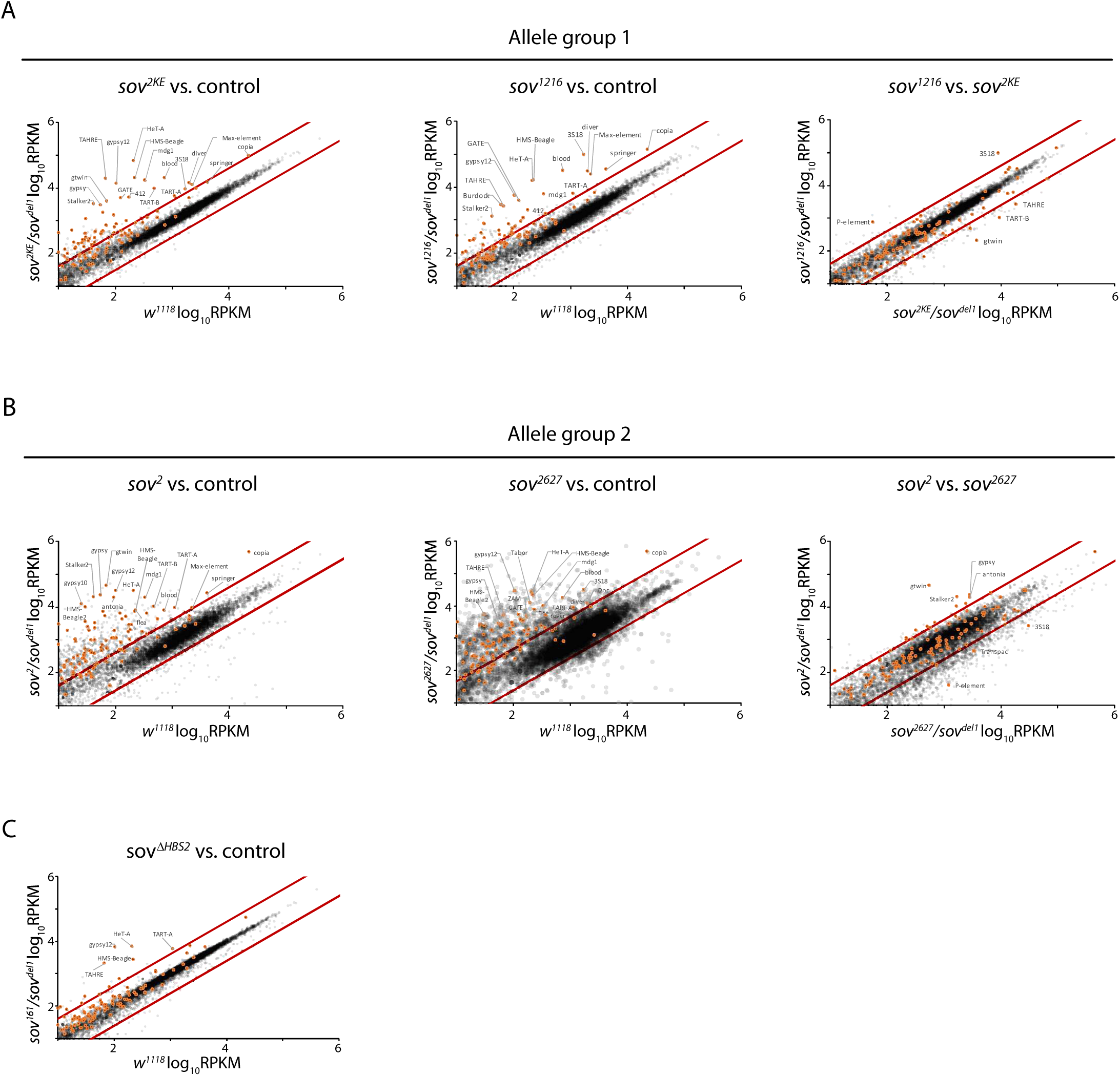
Differential gene expression profiles in various sov mutants, Related to Figure 6. **(A-C)** Scatter plots depict the comparison of steady-state RNA levels (log_10_RPKM) of genes in the indicated *sov* mutant backgrounds compared to each other or to *w¹¹¹* control ovaries. Red lines indicate 2-fold changes. Gray dots represent protein coding genes, transposons are highlighted with orange. **(A)** Allele Group 1 mutants, *sov¹²¹* (left) and *sov^2KE^* (middle), selectively affect transposon repression without altering host gene expression, compared to the *w¹¹¹* control. The highly similar effect of these two alleles on transposon suppression is revealed by comparing their expression profiles (right). **(B)** Global upregulation of host gene and transposon expression in allele Group 2 mutants, *sov²* (left) and *sov² ²* (middle), compared to the *w¹¹¹* control. The effects of the two alleles on global gene expression and transposon suppression are highly similar, as demonstrated by the direct comparison of their effect on gene expression profiles (right). **(C)** Deletion of HBS2 does not alter global gene expression, as demonstrated by steady-state RNA levels in *sov*^Δ*HBS*^^2^*/sov^del1^* ovaries compared to the *w¹¹¹* control. Expression of three telomeric transposons is upregulated.

## SUPPLEMENTAL INFORMATION

TableS1. The area covered by the ligth induced condensates in the optoDroplet assay, related to Figure2D.

TableS2. Primer sequences used for RT-qPCR and ChIP-qPCR used in this study, related to Materials and methods.

TableS3. Mass spectrometry data of the Sov(337–521)-TurboID proximitome, related to Figure5A.

TableS4. Confidence metrics from the pairwise interaction screen between the Sov(337–521) fragment and selected candidate proteins or protein fragments, related to Figure5B.

TableS5. Repair template and gRNA target site sequences used to generate *Drosophila sov* alleles, related to Figure6A.

TableS6. Female fertility tests and phenotype categories of *sov* mutant allelic combinations, related to Figure6B.

TableS7. Normalized RNA-seq read counts generated by DESeq2 from *sov* mutant ovaries, related to Figure6D and FigureS6.

TableS8. *Drosophila* stock used in this study, related to Materials and methods.

MovieS1. A movie sequence showing control nuclei and nuclei of Cry2:mCherry-tagged Sov fragments undergoing droplet formation after light-induced clustering in HEK293T cells.

Scale bars represent 10 µm.

## ACKNOWLEDGEMENTS

We thank the Developmental Studies Hybridoma Bank, the Bloomington *Drosophila* Stock Center, the Vienna *Drosophila* Resource Center and Shelby Blythe for antibodies and fly stocks. We thank the Cellular Imaging Laboratory (CILAB) and the Laboratory of Proteomics at HUN-REN Biological Research Centre, Szeged, for technical support. We are grateful to Margit Szatmári Ugrainé, Anna Rehák and Dorottya Csendes for technical assistance.

